# Sources of outdoor air pollution exposure and child brain network development across the United States

**DOI:** 10.1101/2025.09.23.677862

**Authors:** Katherine L. Bottenhorn, Kirthana Sukumaran, Jordan D. Corbett, Alethea V. de Jesus, Carlos Cardenas-Iniguez, Rima Habre, Daniel A. Hackman, Joel Schwartz, Jiu-Chiuan Chen, Megan M. Herting

## Abstract

Ambient fine particulate matter (PM_2.5_) pollution is a heterogeneous mixture of chemicals with documented neurotoxic effects. Developmental neuroimaging literature has linked childhood PM_2.5_ exposure to alterations in brain morphology, microarchitecture, and function, with implications for cognition and psychopathology. However, the extant literature remains largely cross-sectional and often considers PM_2.5_ a single pollutant, rather than a heterogeneous mixture of chemicals from different sources. This work addresses these gaps by leveraging estimates of exposure to six PM_2.5_ sources derived from positive matrix factorization, and longitudinal neuroimaging data from a large, geographically-diverse sample of Adolescent Brain Cognitive Development Study youth (*N* = 6,291) from across the United States (U.S.). To identify exposure-related differences in brain function and assess their geographical generalizability, we used a predictive modeling approach to assess both differences in functional brain network connectivity during childhood (9-11 years of age) and changes in functional brain network connectivity during the transition to adolescence (9-13 years of age) related to PM_2.5_ exposure. Childhood PM_2.5_ exposure from traffic emissions and industrial/residual fuel burning were linked to mixed patterns of both stronger and weaker connectivity of sensorimotor networks at ages 9-11 years. Conversely, childhood exposures to secondary pollutants (i.e., ammonium sulfates, nitrates) were linked to largely stronger connectivity of brain networks underlying higher-order cognition that decreased over the following two years. However, these patterns of exposure-related functional connectivity identified in youth across the U.S. better represented youth living in the northeast as compared to youth living in the west. Altogether, this work provides insights into the neurotoxicity of outdoor air pollution exposure in developing sensory and motor systems and potential for biomarkers of eventual psychopathology.

## Introduction

Ambient fine particulate matter (<2.5µm in diameter; PM_2.5_) air pollution is among the greatest threats to human health due to its near ubiquity and widespread effects (Costa et al., 2019; Genc et al., 2012). Moreover, PM_2.5_ may be of particular risk to brain health in children (Brumberg et al., 2021), whose higher respiratory rates, levels of physical activity, and time spent outdoors increases their exposure to outdoor PM_2.5_ compared to adults (Bateson & Schwartz, 2007). Beyond increased exposure, children are likely susceptible to longer-term consequences of PM_2.5_ due to their smaller physical size (thus, greater relative dosage) and ongoing development (Gasana et al., 2012). Yet, PM_2.5_ is a complicated mixture of chemicals that are emitted from a host of anthropogenic and non-anthropogenic sources that vary geographically (Snider et al., 2016), and it is unclear which sources cause the most harm to human health. As such, studying differential effects of PM_2.5_ sources on child brain health may provide more accurate and interpretable insight for scientists, policymakers, and parents.

On the cusp of adolescence, ongoing neurodevelopmental changes provide a unique window of susceptibility for outdoor PM_2.5_ to influence brain structure and function, with potential lifespan consequences for cognition and mental health (Herting et al., 2024). In adults, pollution-related neurotoxicity has been linked to increased mental health problems (Braithwaite et al., 2019; Heo et al., 2021; Liu et al., 2021; Qiu et al., 2023) and poorer cognitive performance (Allen et al., 2017; Clifford et al., 2016), but findings regarding exposure-related differences in child and adolescent behavior and cognition are mixed (Campbell et al., 2023; Guxens et al., 2018; Guxens & Sunyer, 2012; Sukumaran et al., 2024), necessitating further study. Emerging human neuroimaging research on the relationships between PM_2.5_ and brain development has focused on macroscale estimates of brain morphology (i.e., global and regional brain volume, cortical thickness (Cserbik et al., 2020; Guxens et al., 2018; Margolis et al., 2022; Miller et al., 2022; Mortamais et al., 2019; Peterson et al., 2022), and, to a lesser extent, microscale estimates (e.g., white matter integrity, gray matter microstructure; (Bottenhorn et al., 2024; Burnor et al., 2021; Lubczyńska et al., 2020; Peterson et al., 2022; Sukumaran et al., 2023)). However, measures of functional brain organization (e.g., connectivity, from resting-state fMRI) have received comparably little attention (Morrel et al., 2025).

Resting state functional connectivity (rsFC) is estimated from functional magnetic resonance imaging (fMRI) data collected while individuals rest, without external stimuli or task paradigms, and is thought to reflect the intrinsic connectivity of large-scale brain networks (collectively, the “connectome”) (Behrens & Sporns, 2012). These networks continue to develop throughout adolescence (Nielsen et al., 2019), support cognition and behavior (Marek et al., 2015; Siegel et al., 2017; Zhu & Qiu, 2022), exhibit transdiagnostic associations with psychopathology (i.e., the triple network model; (V. Menon, 2011)), and are likely vulnerable to neurotoxic impacts of PM_2.5_ exposure. Cross-sectional studies have linked air pollution exposure in the first year of life to pre-adolescent rsFC (i.e., at ages 9-12 years) between default mode and a task positive network, but found that later childhood exposure was unrelated to pre-adolescent rsFC (Pérez-Crespo et al., 2022). Further, childhood exposure to vehicle exhaust was linked to cross-sectional differences in default mode network connectivity and motor response times (Pujol et al., 2016). Only two longitudinal studies have assessed exposure-related rsFC differences, investigating triple network model differences with childhood exposure to PM_2.5_, nitrogen dioxide, and ozone (Cotter et al., 2023; Zundel et al., 2024). This work linked greater PM_2.5_ exposure to increasing default mode network connectivity with both fronto-parietal and salience networks with from 9 to 13 years of age, but these connections weakened with age in youth with lower exposures (Cotter et al., 2023; Zundel et al., 2024). However, no work to date has attempted to disentangle exposure-related differences in functional brain development by investigating multiple sources of PM_2.5_.

Thus, the goal of the current study was to identify childhood differences and longitudinal changes in rsFC patterns related to sources of PM_2.5_ exposure at 9-11 years of age. To do so, we leveraged multi-site, longitudinal neuroimaging and exposure data from the Adolescent Brain Cognitive Development Study^SM^ (ABCD Study®), using connectome-based predictive modeling (CPM; (Shen et al., 2017)) to identify exposure-related patterns of rsFC by PM_2.5_ source during childhood (i.e., at ages 9-11 years) and the transition to adolescence (i.e., over the following two years). Exposure was estimated for six previously established sources of PM_2.5_ (Figure 1A): crustal materials, ammonium sulfates, biomass burning, traffic emissions, ammonium nitrates, and industrial/residual fuel (Sukumaran et al., 2024). Connectivity was estimated between 12 functional networks (Figure 1B): auditory, somatomotor hand and mouth, visual, dorsal and ventral attention, retrosplenial temporal, cingulo-parietal, cingulo-opercular, fronto-parietal, salience, and default mode networks. We did this iteratively, training predictive models with data from all but one ABCD Study site in each iteration and testing said models on data from the held-out site to estimate geographical heterogeneity and generalizability of the identified exposure-related rsFC patterns across the United States. First, we investigated exposure-related cross-sectional *differences* in rsFC at ages 9-11 years with respect to these six sources of ambient PM_2.5_ (Figure 1C, #1). Then, we investigated exposure-related longitudinal *changes* in rsFC at ages 9-11 years with respect to these six sources of ambient PM_2.5_ over the following 2 years (i.e., from 9 to 13 years of age) (Figure 1C, #2). Altogether, this work revealed more detailed patterns of exposure-related rsFC during the transition to adolescence, proposing nuanced roles of different PM_2.5_ sources.

**Figure 1.**
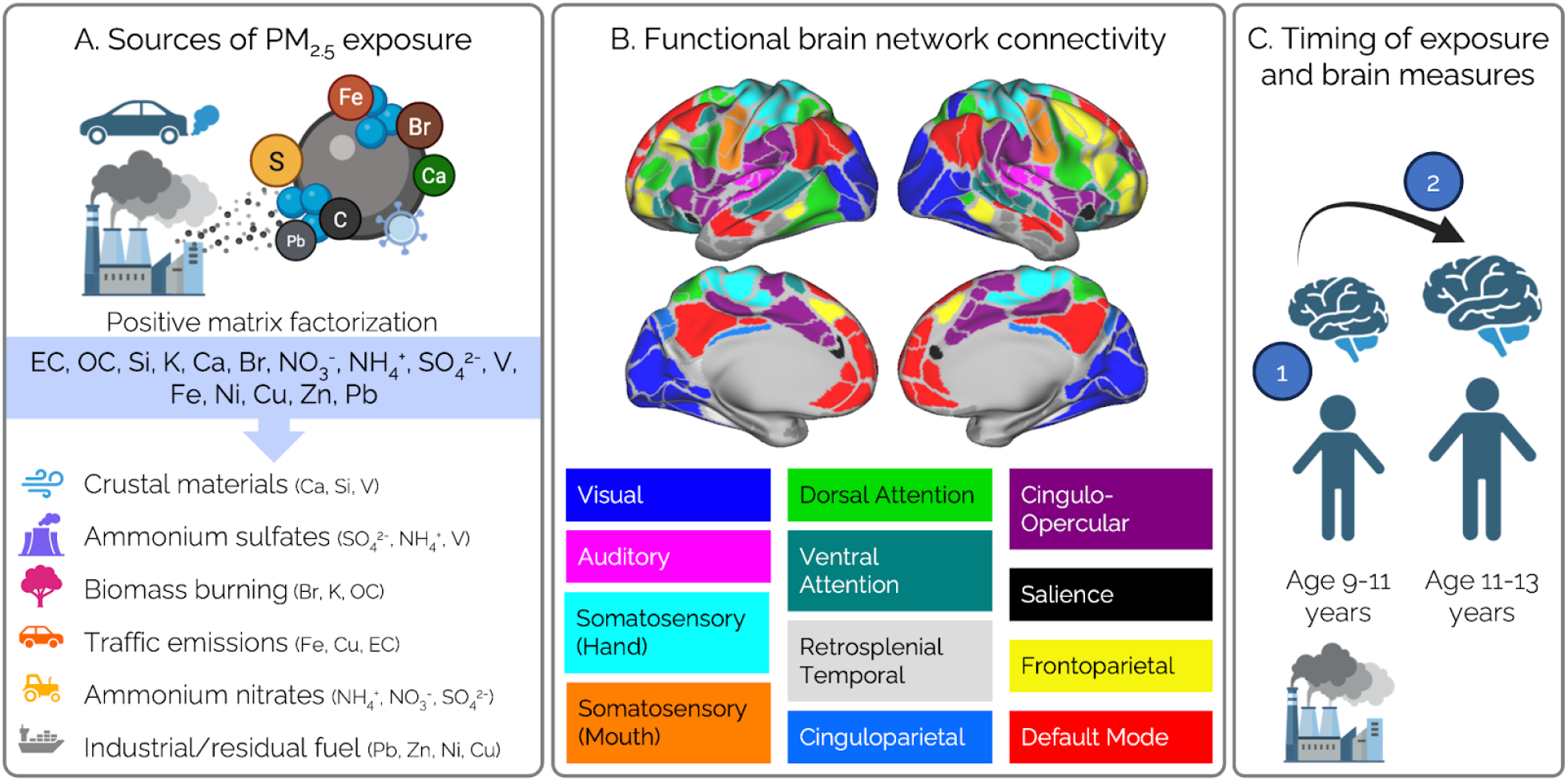
Linking sources of child PM_2.5_ exposure to functional brain network connectivity both at ages 9-11 years and longitudinally during the transition to adolescence. (A) Building on the previously identified six common sources of PM_2.5_ across the United States via positive matrix factorization of 15 PM_2.5_ components (Sukumaran et al., 2024), this work addresses source-related exposure differences in resting-state functional connectivity. (B) Functional connectivity was assessed between 12 large-scale functional brain networks, involved in sensorimotor processing (left column: auditory, visual, somatosensory–hand & mouth), attention and memory (middle column: dorsal and ventral attention, retrosplenial temporal, cingulo-parietal), and higher-order functions (right column: fronto-parietal, salience, default mode, cingulo-opercular). (C) We assessed associations between exposure to PM_2.5_ sources in childhood (i.e., at 9-11 years of age) and (1) functional connectivity at 9-11 years of age (i.e., cross-sectionally) as well as (2) changes in functional connectivity over the following two years, during the transition to adolescence (i.e., longitudinally, from 9 to 13 years of age).

## Methods

### Participants

As a part of the ongoing Adolescent Brain and Cognitive Development (ABCD) Study, longitudinal data were obtained from the annual 5.1 data release (https://dx.doi.org/10.15154/z563-zd24). The ABCD Study enrolled over 11,000 children 9 to 10 years of age in a 10-year longitudinal study between October 2016 and October 2018 (Garavan et al., 2018; Volkow et al., 2018). Participants were recruited at 21 study sites across the U.S. from elementary schools (private, public, and charter schools) and twin registries in a design intended to assemble a nationally-representative sample, with respect to sociodemographic diversity (Garavan et al., 2018). The institutional review board and human research protections programs at the University of California San Diego approved all experimental and consent procedures, as did the local institutional review board of each study site. Each participant provided written assent in addition to their legal guardian’s written agreement to participate and consent to participate, themselves. Exclusion criteria for the ABCD study included lack of English proficiency, intellectual or medical limitations, severe sensory sensitivities, neurological disorders, and inability to complete an MRI scan (Palmer et al., 2022).

Here, we use a subset of ABCD Study data from two time points: collected at ages 9-11 years and 11-13 years, including residential estimates of PM_2.5_ exposure, magnetic resonance imaging (MRI), measures of participants’ sex at birth, socio-demographics, and pubertal development. Measures of participants’ sex, demographics, and PM_2.5_ estimates were collected at ages 9-11 years. Participant selection for our analyses is outlined in Supplementary Figure 1. Participants were only included in the following analyses who: had no incidental neurological findings evident in their scans, passed the ABCD imaging or FreeSurfer quality control procedures (Hagler Jr et al., 2019), and had sufficient quality resting-state fMRI data (i.e., framewise displacement <0.5 mm average for head motion, at least 10 minutes of low-motion data). For analyses including rsFC *changes*, our sample was narrowed to participants with fMRI data collected on Siemens MRI scanners, due to software updates between waves of data collection in General Electric and Philips MRI scanners. We also only included participants if their 2-year follow-up visit before March 1, 2020, to avoid potentially confounding effects of differential COVID-19 experiences on exposure or neurodevelopmental measures. We, additionally, omitted data from one of the 21 study sites due to hardware issues identified during baseline data collection. The final sample characteristics for the current study are described in Table 1.

**Table 1.**
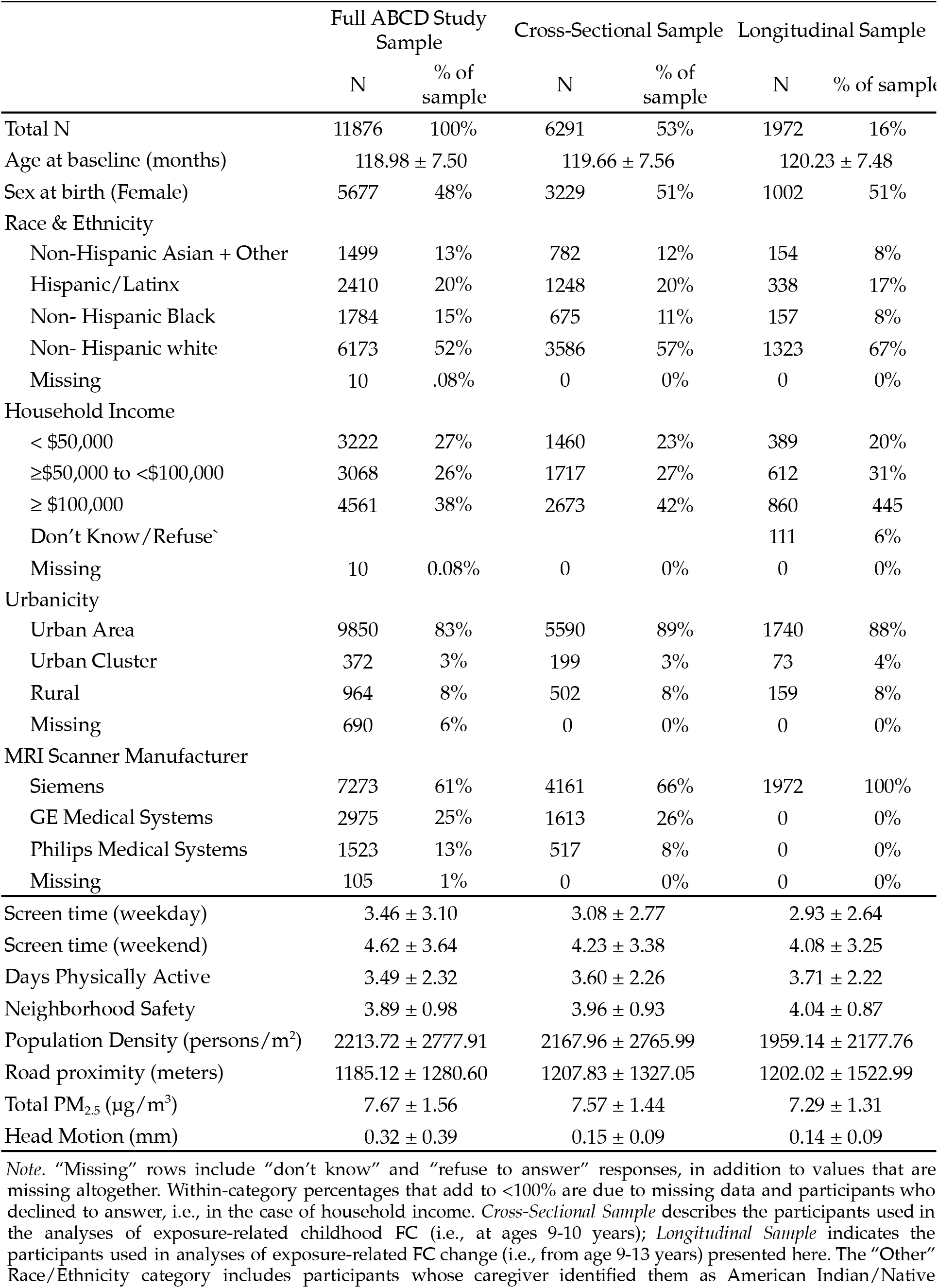
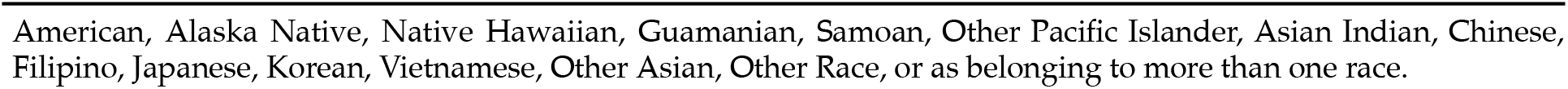
Sample Demographics.

|  | Full ABCD Study Sample |  | Cross-Sectional Sample |  | Longitudinal Sample |  |
| --- | --- | --- | --- | --- | --- | --- |
|  | N | % of sample | N | % of sample | N | % of sample |
| Total N | 11876 | 100% | 6291 | 53% | 1972 | 16% |
| Age at baseline (months) | 118.98 ± 7.50 |  | 119.66 ± 7.56 |  | 120.23 ± 7.48 |  |
| Sex at birth (Female) | 5677 | 48% | 3229 | 51% | 1002 | 51% |
| Race & Ethnicity |  |  |  |  |  |  |
| Non-Hispanic Asian + Other | 1499 | 13% | 782 | 12% | 154 | 8% |
| Hispanic/Latinx | 2410 | 20% | 1248 | 20% | 338 | 17% |
| Non- Hispanic Black | 1784 | 15% | 675 | 11% | 157 | 8% |
| Non- Hispanic white | 6173 | 52% | 3586 | 57% | 1323 | 67% |
| Missing | 10 | .08% | 0 | 0% | 0 | 0% |
| Household Income |  |  |  |  |  |  |
| < \$50,000 | 3222 | 27% | 1460 | 23% | 389 | 20% |
| ≥\$50,000 to <\$100,000 | 3068 | 26% | 1717 | 27% | 612 | 31% |
| ≥ \$100,000 | 4561 | 38% | 2673 | 42% | 860 | 44% |
| Don't Know/Refuse` |  |  |  |  | 111 | 6% |
| Missing | 10 | 0.08% | 0 | 0% | 0 | 0% |
| Urbanicity |  |  |  |  |  |  |
| Urban Area | 9850 | 83% | 5590 | 89% | 1740 | 88% |
| Urban Cluster | 372 | 3% | 199 | 3% | 73 | 4% |
| Rural | 964 | 8% | 502 | 8% | 159 | 8% |
| Missing | 690 | 6% | 0 | 0% | 0 | 0% |
| MRI Scanner Manufacturer |  |  |  |  |  |  |
| Siemens | 7273 | 61% | 4161 | 66% | 1972 | 100% |
| GE Medical Systems | 2975 | 25% | 1613 | 26% | 0 | 0% |
| Philips Medical Systems | 1523 | 13% | 517 | 8% | 0 | 0% |
| Missing | 105 | 1% | 0 | 0% | 0 | 0% |
| Screen time (weekday) | 3.46 ± 3.10 |  | 3.08 ± 2.77 |  | 2.93 ± 2.64 |  |
| Screen time (weekend) | 4.62 ± 3.64 |  | 4.23 ± 3.38 |  | 4.08 ± 3.25 |  |
| Days Physically Active | 3.49 ± 2.32 |  | 3.60 ± 2.26 |  | 3.71 ± 2.22 |  |
| Neighborhood Safety | 3.89 ± 0.98 |  | 3.96 ± 0.93 |  | 4.04 ± 0.87 |  |
| Population Density (persons/m<sup>2</sup>) | 2213.72 ± 2777.91 |  | 2167.96 ± 2765.99 |  | 1959.14 ± 2177.76 |  |
| Road proximity (meters) | 1185.12 ± 1280.60 |  | 1207.83 ± 1327.05 |  | 1202.02 ± 1522.99 |  |
| Total PM<sub>2.5</sub> (µg/m<sup>3</sup>) | 7.67 ± 1.56 |  | 7.57 ± 1.44 |  | 7.29 ± 1.31 |  |
| Head Motion (mm) | 0.32 ± 0.39 |  | 0.15 ± 0.09 |  | 0.14 ± 0.09 |  |

*Note.* "Missing" rows include "don't know" and "refuse to answer" responses, in addition to values that are missing altogether. Within-category percentages that add to <100% are due to missing data and participants who declined to answer, i.e., in the case of household income. *Cross-Sectional Sample* describes the participants used in the analyses of exposure-related childhood FC (i.e., at ages 9-10 years); *Longitudinal Sample* indicates the participants used in analyses of exposure-related FC change (i.e., from age 9-13 years) presented here. The "Other" Race/Ethnicity category includes participants whose caregiver identified them as American Indian/Native
American, Alaska Native, Native Hawaiian, Guamanian, Samoan, Other Pacific Islander, Asian Indian, Chinese, Filipino, Japanese, Korean, Vietnamese, Other Asian, Other Race, or as belonging to more than one race.

### Demographic Data, Confounders, & Precision Variables

Potential confounders and precision variables were used to create a directed acyclic graph (DAG; Supplementary Figure 2), based on theoretical knowledge of factors associated with brain development and air pollution exposure (Bottenhorn et al., 2024; Sukumaran et al., 2024). Because pollution levels are higher in minority communities and those from disadvantaged social status backgrounds, sociodemographic variables are included (Hajat, Hsia, and O’Neill 2015). Scanner (i.e., MRI) manufacturer (Siemens, Philips, GE) is included in the cross-sectional analyses to account for differences in image acquisition, but only data collected on Siemens scanners is used for longitudinal analyses due to inconsistent software updates on Philips and GE MRIs (see ABCD Study 5.1 release notes). From this DAG, a minimally sufficient set of covariates were identified, which represent the smallest group of variables that can account for all paths by which the entire variable set may impact air pollution exposure and cortical microstructure development. The minimally sufficient set that was used as covariates in the analyses presented here included race/ethnicity (non-Hispanic *white, non-Hispanic Black, Hispanic/Latinx, non-Hispanic Asian*, or *Other*), combined household income in USD (*≥$100K, ≤$50K and <$100K, <$50K*, or *Don’t Know/Refuse to Answer*), caregiver-reported perceived neighborhood safety based on a survey modified from PhenX (NSC) (Echeverria et al., 2004; Mujahid et al., 2007), and the following information about the child’s primary residence based on geospatial estimates: the population density (Center For International Earth Science Information Network-CIESIN-Columbia University, 2018; Fan et al., 2021), urbanicity (*US Census Track Urban Classification*), and distance to major roadways (meters) (Fan et al., 2021). Additional variables were included as key sociodemographic and behavioral precision variables that might influence outdoor exposure levels, including the child’s age and sex assigned at birth (*male* or *female*) as well as average daily screen time (in hours, separate for weekday and weekend) and physical activity (number of days child was physically active in the week prior). Additional precision variables included the manufacturer of the MRI on which each participant’s data were collected to account for differences in both scanner hardware and software (Ciric et al. 2018), handedness (*right, left*, or *mixed*) to account for potential brain differences in laterality, as well as average head motion (framewise displacement, in mm) during fMRI scans.

**Figure 2.**
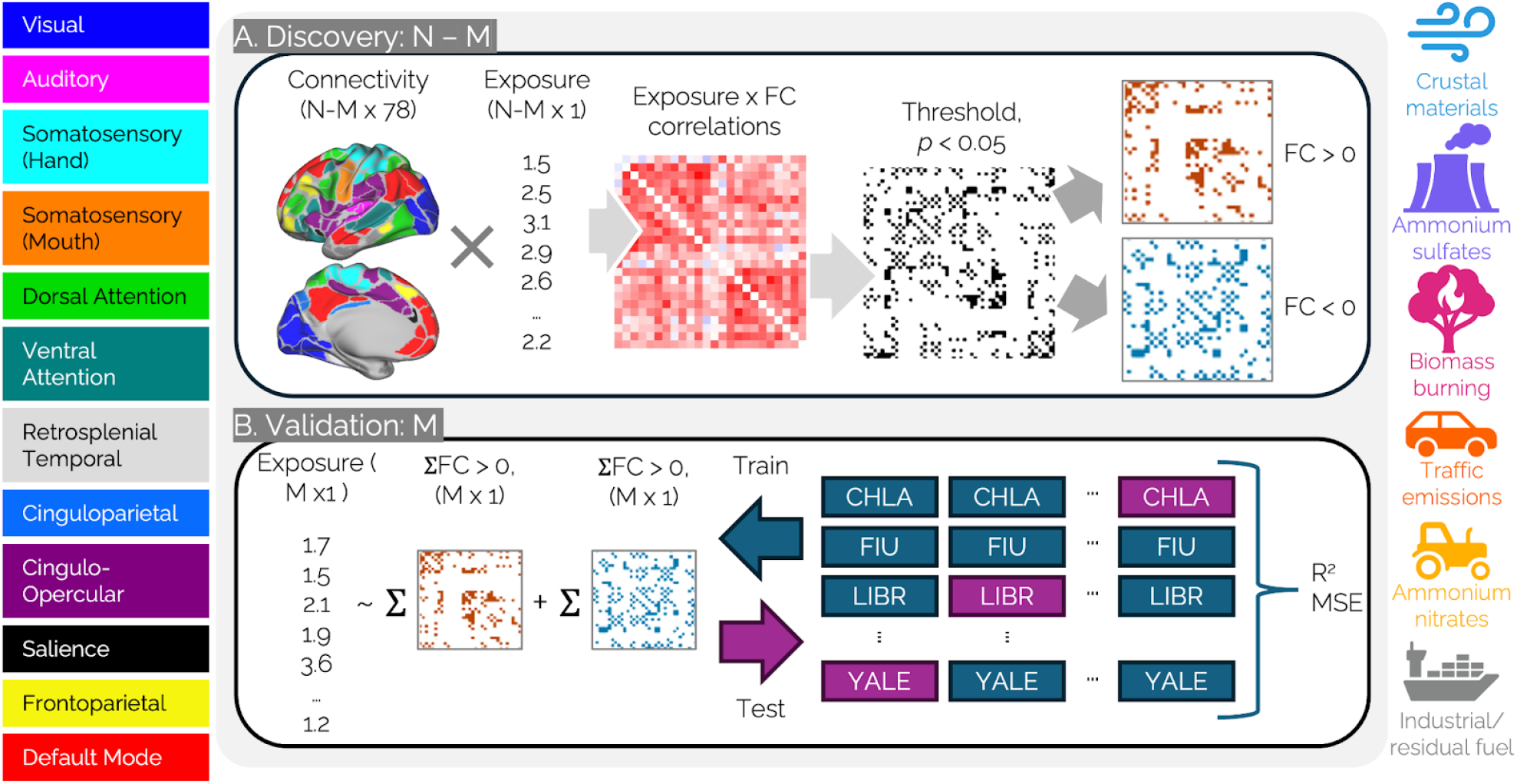
Predictive connectomics schematic. A) Identifying source-specific PM_2.5_ exposure-related connectivity between large-scale functional brain networks and B) testing geographical generalizability of results. Functional connectivity of 12 large-scale brain networks (color-coded, left) and six source-specific PM_2.5_ exposure estimates (color-coded icons, right) were first used in the (A) Discovery sample (participants from all sites, *N*, minus one, *M*) to identify PM_2.5_ source-related connectivity by correlating each participant’s FC and exposure, cross-sectionally at ages 9-10 years (Analysis 1) and longitudinal changes from ages 9-13 years old (Analysis 2). Exposure x FC correlations were thresholded at *p* < 0.05, from which connections positively associated with exposure (i.e., *FC* > 0) and negatively associated with exposure (*FC* < 0) were separated. (B) Next, in the Validation sample, the generalizability of findings across geographically distinct ABCD Study sites was assessed. First, training a linear regression model was trained on data from all sites, except one site (*M*), to predict exposure estimates from the sum of positively- and negatively-related FC estimated coefficients for each functional connection. Then,the resulting regression model was tested in participants from the remaining, held-out site (*M*. This process was repeated for each source of PM_2.5_. The six common sources of PM_2.5_ include crustal materials (e.g., wind-blown dust) in blue, ammonium sulfates (e.g., from coal-burning power plants) in purple, biomass burning (e.g., wildfires) in pink, traffic emissions (i.e., both tailpipe and non-tailpipe emissions) in orange, ammonium nitrates (e.g., from crop fertilizers) in yellow, and industrial/ residual fuel burning in gray. Abbreviations: coefficient of variation, *R*^2^; mean squared error, MSE; product-moment correlation coefficient, *r*.

### Neuroimaging Data

#### Resting State fMRI (rs-fMRI): Acquisition and Quality Control

Each scan was collected in accordance with harmonized procedures on Siemens Prisma, Philips, or GE 750 3T MRI scanners. Imaging acquisition protocols specific to the ABCD Study have been described by Casey et al. (2018). Twenty cumulative minutes of resting-state data was collected across two sets of two five-minute acquisition periods, while subjects were instructed to keep their eyes open and fixed on a crosshair. This protocol increases the probability of collecting enough data with low motion per ABCD’s standards (>12.5 minutes of data with framewise displacement (FD) < 0.2 mm) (Power et al. 2014). Resting state scans were acquired using an echo-planar imaging sequence in the axial plane, with the following parameters: TR = 800 ms, TE = 30 ms, flip angle = 90°, voxel size = 2.4 mm^3^, 60 slices. Only images without clinically significant incidental findings (*mrif_score* = 1 or 2) that passed all ABCD quality-control parameters were included in analysis (*imgincl_rsfmri_include* = 1). To ensure sufficient, high-quality data, we further excluded imaging data from participants with fewer than 10 minutes of low-motion data and with greater than 0.5mm average head motion across the fMRI scans.

#### rs-fMRI Processing: Cross-sectional and Longitudinal Changes in Functional Connectivity

Image processing steps have been previously described by Hagler and colleagues (2019). Twelve canonical networks were functionally defined using resting-state rsFC patterns according to methods described by Gordon et al. (2016). These included auditory, visual, somatomotor (hand), somatomotor (mouth), dorsal attention, ventral attention, retrosplenial temporal, cinguloparietal, cingulo-opercular, salience, frontoparietal, and default mode networks. Intra-network correlations were calculated by averaging pairwise product-moment correlations for ROIs belonging to that network; inter-network correlations were calculated by averaging pairwise product-moment correlations between ROIs within the first network and ROIs within the second network (Gordon et al. 2016). Using the Fisher transformation, these correlations were then converted to *z*-scores.

Longitudinal changes in rsFC were calculated as “reliable changes”, adapted for use with rsFC, which was calculated from product-moment correlations between BOLD signals from each network, ranging from −1.0 to 1.0, with −1.0 indicating “perfect anticorrelation” and 1.0 indicating perfect correlation. Because correlation and anticorrelation likely reflect the same degree of information transfer, but with different effects (i.e., both activated vs. one activated and the other deactivated), we considered them equivalent “connectivity” between brain networks. Thus, reliable changes in rsFC (ΔFC; Equation 1; (Bottenhorn et al., 2025)) are calculated from the *absolute values* of the *z*-scored correlation between two networks, at each time point (i.e., at ages 9-11 years, *FC*_*T*1_, and at ages 11-13 years, *FC*_*T*2_, as in Eq. 1 below), to reflect the change in “connectedness”, and then scaled by the age difference (i.e., *age*_*T*2_ - *age*_*T*1_ in Eq. 1 below). Thus, analyses using reliable change scores can assess how sources of PM_2.5_ exposure are related to changes in information transfer across the brain, as reflected by rsFC, while accounting for differences in measurement error (e.g., head motion) between time points.

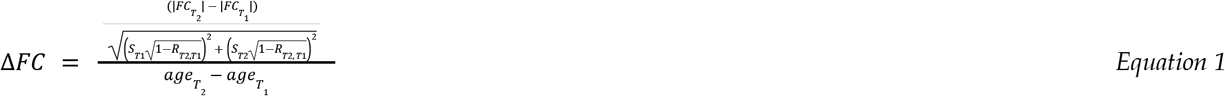

Where *ΔFC* represents a reliable change in rsFC, *FC*_*T*1_ represents rsFC at 9-11 years of age, *FC*_*T*2_ represents rsFC at 11-13 years of age, *s*_*T*1_ represents variance in rsFC at 9-11 years of age across the sample, *s*_*T*2_ represents variance in rsFC at 11-13 years of age across the sample, *R*_*T2, T1*_ represents the correlation between rsFC at ages 9-11 and 11-13 years of age across the sample, *age*_*T1*_ represents participant age at baseline data collection, and *age*_*T2*_ represents participant age at 2-year follow-up data collection.

### Ambient Air Pollution Estimates

Annual ambient air pollution concentrations for fine particulate matter (PM_2.5_) and from fifteen PM_2.5_ components (i.e., nickel, Ni; iron, Fe; bromine, Br; vanadium, V; potassium, K; lead, Pb; zinc, Zn; copper, Cu; calcium, Ca; silicon, Si; elemental carbon, EC; sulfates, SO_4_^2-^; nitrates, NO_3_^-^; ammonium, NH_4_^+^; and organic carbon, OC) were assigned to primary residential addresses of each child (Cardenas-Iniguez et al., 2024; Fan et al., 2021). Briefly, daily PM_2.5_ estimates were derived at a 1-km^2^ resolution whereas annual PM_2.5_ component estimates were estimated at the 50 meter resolution using hybrid spatiotemporal models, which utilize satellite-based aerosol optical depth models, chemical transport models, and land-use regression (Amini et al., 2023; Di et al., 2019). Performance of these spatiotemporal models are reported in Supplementary Table 1. These estimates were then averaged over the 2016 calendar year, corresponding with the inception of the baseline ABCD Study visits (October 2016-2018). These concentrations were then assigned to the primary residential address at the baseline study visit when children were aged 9-11 years (Supplementary Table 2). As previously published, we used positive matrix factorization (PMF; United States Environmental Protection Agency, EPA v5.0) alongside participant level concentrations of the fifteen PM_2.5_ components, to identify six common sources of PM_2.5_ across all ABCD Study participants from the 21 study sites (Supplementary Figure 3; (Sukumaran et al., 2024)). These six sources corresponded with crustal materials (i.e., soil, dust; driven by V, Si, Ca), ammonium sulfates (e.g., from coal burning; driven by SO_4_^2-^, NH_4_^+^, V), biomass burning (e.g., from wildfires; driven by Br, K, OC), traffic (i.e., both tailpipe and non-tailpipe emissions; driven by Fe, Cu, EC), ammonium nitrates (e.g., from agricultural fertilizers and NO_2_ sources; driven by NH_4_^+^, NO_3_^-^, SO_4_^2-^), and industrial emissions/residual fuel burning (e.g., from heavy fuel combustion; driven by Pb, Zn, Ni, Cu). Individual-level source apportionment contributions for each of these six commonly identified sources were then utilized as exposure estimates in the following analyses (Supplementary Table 3). Product moment correlations of individual-level estimates from each of the six common sources are reported in Supplementary Figure 4.

**Figure 3.**
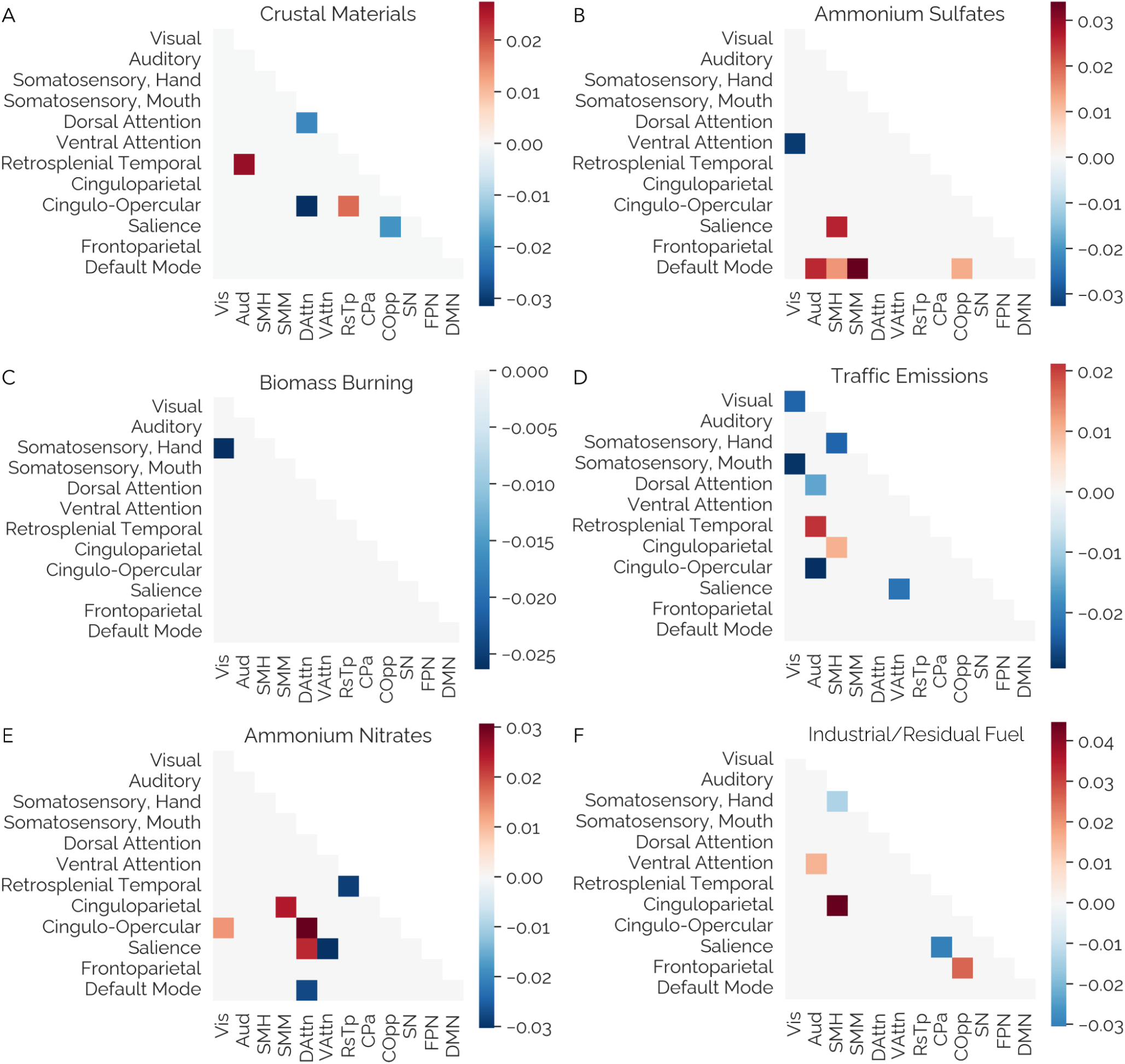
Exposure-related differences in functional connectivity at ages 9-10 years-old. Patterns of functional connectivity associated with exposure to PM_2.5_ from A) crustal materials, B) ammonium sulfates, C) biomass burning, D) traffic emissions, E) ammonium nitrates, and F) industrial/residual fuel (averaged across CV loops and thresholded at |*r*_*S*_| > 0.01). Lower triangles of connectivity matrices per PM_2.5_ source exhibit exposure-related functional network connectivity. Colors reflect correlations between rsFC and exposure estimates. Positive values indicate relative exposure-related strengthening of connectivity between two networks, while negative values indicate exposure-related weakening of connectivity. Network abbreviations: visual, Vis; auditory, Aud; somatosensory hand, SMH; somatosensory mouth, SMM; dorsal attention, DAttn; ventral attention, VAttn; retrosplenial temporal, RsTp; cinguloparietal, CPa; cingulo-opercular, COpp; salience, SN; frontoparietal, FPN; default mode, DMN.

**Figure 4.**
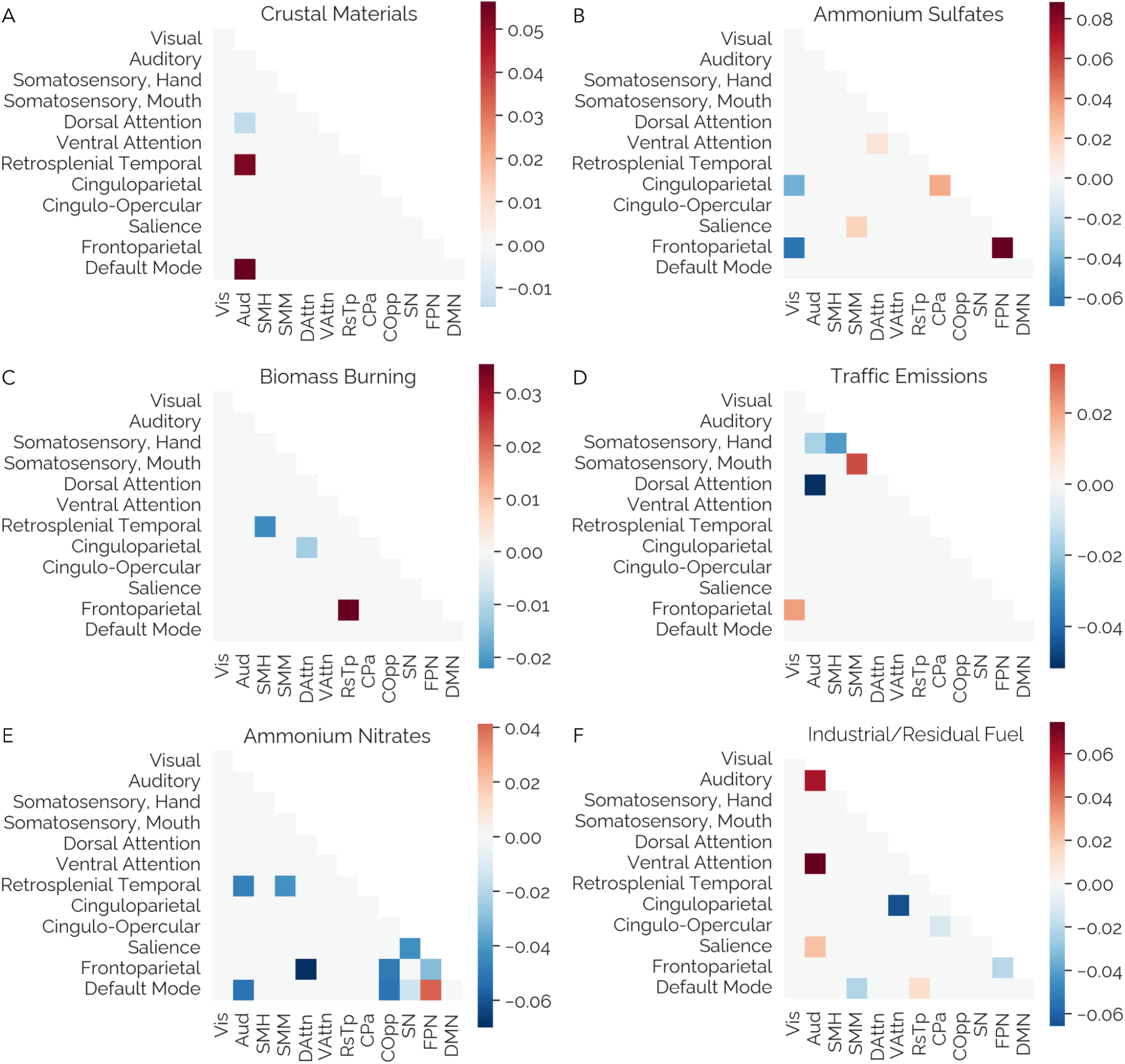
Exposure-related, longitudinal changes in functional connectivity (ΔFC) from ages 9-13 years. Changes in functional connectivity associated with exposure to PM_2.5_ from A) crustal materials, B) ammonium sulfates, C) biomass burning, D) traffic emissions, E) ammonium nitrates, and F) industrial/residual fuel (averaged across CV loops and thresholded at |*r*_*S*_| > 0.01). Colors reflect pairwise product-moment correlations between **ΔFC** and exposure estimates, inverse weighted by the average RMSE (i.e., between predicted and actual exposure estimates in participants from the held-out site) across the leave-one-site-out cross-validation. Positive values indicate exposure-related strengthening of connectivity between two networks, while negative values indicate exposure-related weakening of connectivity. Average RMSE values can be found in Table 3. Network abbreviations: visual, Vis; auditory, Aud; somatosensory hand, SMH; somatosensory mouth, SMM; dorsal attention, DAttn; ventral attention, VAttn; retrosplenial temporal, RsTp; cinguloparietal, CPa; cingulo-opercular, COpp; salience, SN; frontoparietal, FPN; default mode, DMN.

### Predictive Connectomics

To identify exposure-related rsFC patterns for multiple sources of PM_2.5_, we used connectome-based predictive modeling (CPM; (Shen et al., 2017). CPM leverages network science and machine learning principles for a reliable, interpretable assessment of rsFC (Finn & Rosenberg, 2021; Mantwill et al., 2022; Shen et al., 2017). Specifically, using CPM, we first assessed cross-sectional *differences* in functional network connectivity at ages 9-11 years old related to exposure to each source of PM_2.5_ (Analysis 1). Then, CPM was used to determine whether source-related exposure influenced longitudinal *changes* in functional network connectivity between 9 and 13 years of age (Analysis 2).

In each analysis, CPM was used to identify rsFC related to each PM_2.5_ source separately, in a leave-one-site-out cross-validation (CV) framework. Within each CV loop (i.e., in all ABCD Study sites except one), we first regressed out the minimally sufficient covariate set from rsFC and exposure estimates (including data collection site as a fixed effect). Then, exposure-related rsFC was identified in individuals from 19 of the 20 sites (Figure 2A) by performing connection-wise *F*-tests for connectivity related to pollution source loadings. Connections significantly related to exposure (at *p* < 0.05) were separated into positively-related rsFC (*r* > 0) and negatively-related rsFC (*r* < 0). For each participant, all positively-related connections (*FC*_+_ in *Equations 2a-b*) were summed and all negatively-related connections (*FC*_-_ in *Equations 2a-b*) were summed, separately, creating vectors of network strength positively- and negatively-related to exposure. These vectors were then used as independent variables in a linear regression (*Equation 2*) predicting exposure to each source (*Exposure*_*S*_ in *Equations 2a-b*). Second, the fit of this linear model of exposure-related rsFC was assessed using data from the held-out site (Figure 2B). By using these equations to predict exposure from rsFC, we were able to quantify heterogeneity in model fit across study sites by calculating the root mean squared error (RMSE) and coefficient of variation (*R*^2^) from predicted and actual exposure values in both the training sites and the held-out test site.

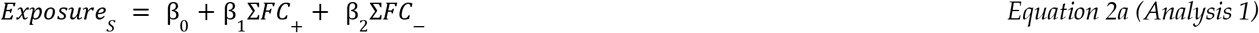

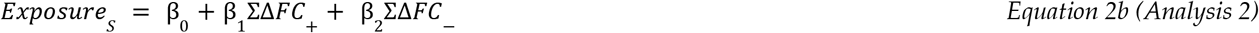

RMSE indicates how accurately the identified rsFC connectivity pattern is able to estimate individual-level exposures, with lower values indicating less error between actual and predicted values and, thus, a better performing model. Additionally, coefficient of determination (*R*^2^) indicates how much of the variability in exposure is explained by the identified rsFC patterns, with higher values indicating greater agreement between the predicted and actual exposure values and, thus, a better performing model. In other words, in the current study a lower RMSE and a higher *R*^2^ indicated greater geographic generalizability between sources of exposure and rsFC patterns.

## Results

Our final sample size for exposure-related *differences* in functional brain connectivity at 9-11 years of age (Analysis 1, cross-sectional sample) was *N* = 6,291 (51% female) and *changes over time* from 9 to 13 years of age (Analysis 2, longitudinal sample) was *N* = 1,972 (50% female) with an average of 23.99 ± 1.63 months between visits (Table 1). Annual average total PM_2.5_ exposure was 7.57 ± 1.44 µg/m^3^ in the cross-sectional sample and 7.29 ± 1.31 µg/m^3^ in the longitudinal subsample.

### Source-related functional connectomics at ages 9-11 years

Across the U.S.-wide sample (*N* = 6,291), connectome-based predictive modeling identified exposure-related patterns of rsFC differences at ages 9-11 years associated with each of the six common sources of PM_2.5_ (Figure 3). Greater crustal material exposure (Figure 3A) was positively related to connectivity of the retrosplenial temporal network (i.e., with auditory and cingulo-opercular networks), but negatively related to connectivity within the dorsal attention network and connectivity between the cingulo-opercular network and both salience and dorsal attention networks. For ammonium sulfates (Figure 3B), exposure was positively related to default mode network connectivity (i.e., with auditory, somatomotor, and cingulo-opercular networks) and salience network connectivity (i.e., with somatomotor-hand networks). Conversely, ammonium sulfate exposure was negatively related to connectivity between the visual and ventral attention networks. Biomass burning exposure (Figure 3C) was negatively related to connectivity between the visual and somatomotor-hand networks. Exposure from traffic emissions (Figure 3D) was positively related to connectivity between retrosplenial temporal and auditory networks, as well as between cinguloparietal and somatomotor-hand networks. Traffic emissions exposure was negatively related to connectivity both within and between visual and somatomotor (hand) networks, in addition to auditory network (i.e., with cingulo-opercular and dorsal attention networks) and ventral attention network (i.e., with salience network) connectivity. Ammonium nitrates (Figure 3E) was positively related to connectivity of the somatomotor-mouth network (i.e., with cingulo-parietal network), the cingulo-opercular network (i.e., with visual and dorsal attention networks), and the dorsal attention network (i.e., with salience and cingulo-opercular networks). Ammonium nitrate exposure was also negatively related to connectivity within the retrosplenial temporal network, as well as between default mode and dorsal attention networks. Interestingly, ammonium nitrates are positively related to salience network connectivity with the dorsal attention network, but negatively related to salience network connectivity with the ventral attention network. Industrial/residual fuel burning (Figure 3F) was positively related to auditory network connectivity (i.e., with the ventral attention network), connectivity between cinguloparietal and somatomotor-hand networks, and between frontoparietal and cingulo-opercular networks. Conversely, industrial/residual fuel exposure was negatively associated with connectivity within the somatomotor-hand network and between cinguloparietal and salience networks. Across sources, traffic emissions were most often related to weaker rsFC, while ammonium sulfates were most often related to stronger rsFC (primarily between the default mode network and sensorimotor networks).

### Source-specific changes in longitudinal functional connectomics from ages 9 to 13 years

Across the smaller U.S. sample of participants with MRI data collected on Siemens scanners (*N* = 1,972), predictive connectomics identified patterns of longitudinal changes in rsFC related to each of the six sources (Figure 4). Exposure to crustal materials (Figure 4A) was related to changes in auditory network connectivity: stronger connectivity with default mode and retrosplenial temporal networks, but weaker connectivity with the dorsal attention network over the two year follow-up. For ammonium sulfates (Figure 4B), greater exposure was related to increases in dorsal-to-ventral attention network connectivity and salience-to-somatomotor network connectivity, in addition to within-network cingulo-parietal and within-network frontoparietal connectivity over time. Conversely, greater ammonium sulfate exposure was related to decreases in visual network connectivity (i.e., with cingulo- and frontoparietal networks) over two years. Greater biomass burning exposure (Figure 4C) was related to increases in connectivity between retrosplenial temporal and frontoparietal networks, but decreases in connectivity between retrosplenial temporal and somatomotor-hand networks, as well as between dorsal attention and cinguloparietal networks over time. Exposure to traffic emissions (Figure 4D) was related to increasing connectivity within the somatomotor-mouth network and between frontoparietal and visual networks, but related to decreasing connectivity within the somatomotor-hand networks, as well as between auditory and both somatomotor-hand and dorsal attention networks over the two year follow-up. Exposure to ammonium nitrates (Figure 4E) was related to increased connectivity between frontoparietal and default mode networks over time. Conversely, greater ammonium nitrate exposure was related to decreases in salience (i.e., within network and with default mode), frontoparietal (i.e., within network and with dorsal attention, cingulo-opercular networks), retrosplenial temporal (i.e., with auditory and somatomotor-mouth networks), and default mode (i.e., with auditory, cingulo-opercular, and salience networks) network connectivity. Exposure to industrial/residual fuel burning (Figure 4F) was related to increases in auditory network connectivity (i.e., with ventral attention, auditory, and salience networks), as well as connectivity between retrosplenial and default mode networks over time. Conversely, industrial/residential fuel exposure was related to connectivity decreases in cinguloparietal (i.e., with ventral attention, cingulo-opercular networks), default mode (i.e., with somatomotor-mouth), and frontoparietal (i.e., within-network) networks. Across sources, ammonium nitrates were most often related to decreases in rsFC (especially in higher-order networks), while ammonium sulfates and industrial/residual fuel burning were most often related to increases in rsFC, although to a lesser extent.

### Exposure-related functional connectivity by network

Finally, we summarize exposure-related rsFC by network (Figure 5). Across sources of PM_2.5_, the visual, dorsal attention, and salience networks most often exhibited weaker connectivity at ages 9-11 years (i.e., cross-sectionally) with greater exposure, while the auditory, somatosensory-hand, cingulo-opercular, and default mode most often exhibited stronger connectivity. The dorsal attention network exhibited the most exposure-related decreases in rsFC across sources from 9-13 years of age (i.e., longitudinally), as did the auditory, cinguloparietal, frontoparietal, and default mode networks. Similarly, the auditory and default mode networks exhibited the most exposure-related increases in rsFC from 9-13 years of age. Thus, auditory, dorsal attention, frontoparietal, and default mode networks may be particularly sensitive to exposure-related neurotoxicity.

**Figure 5.**
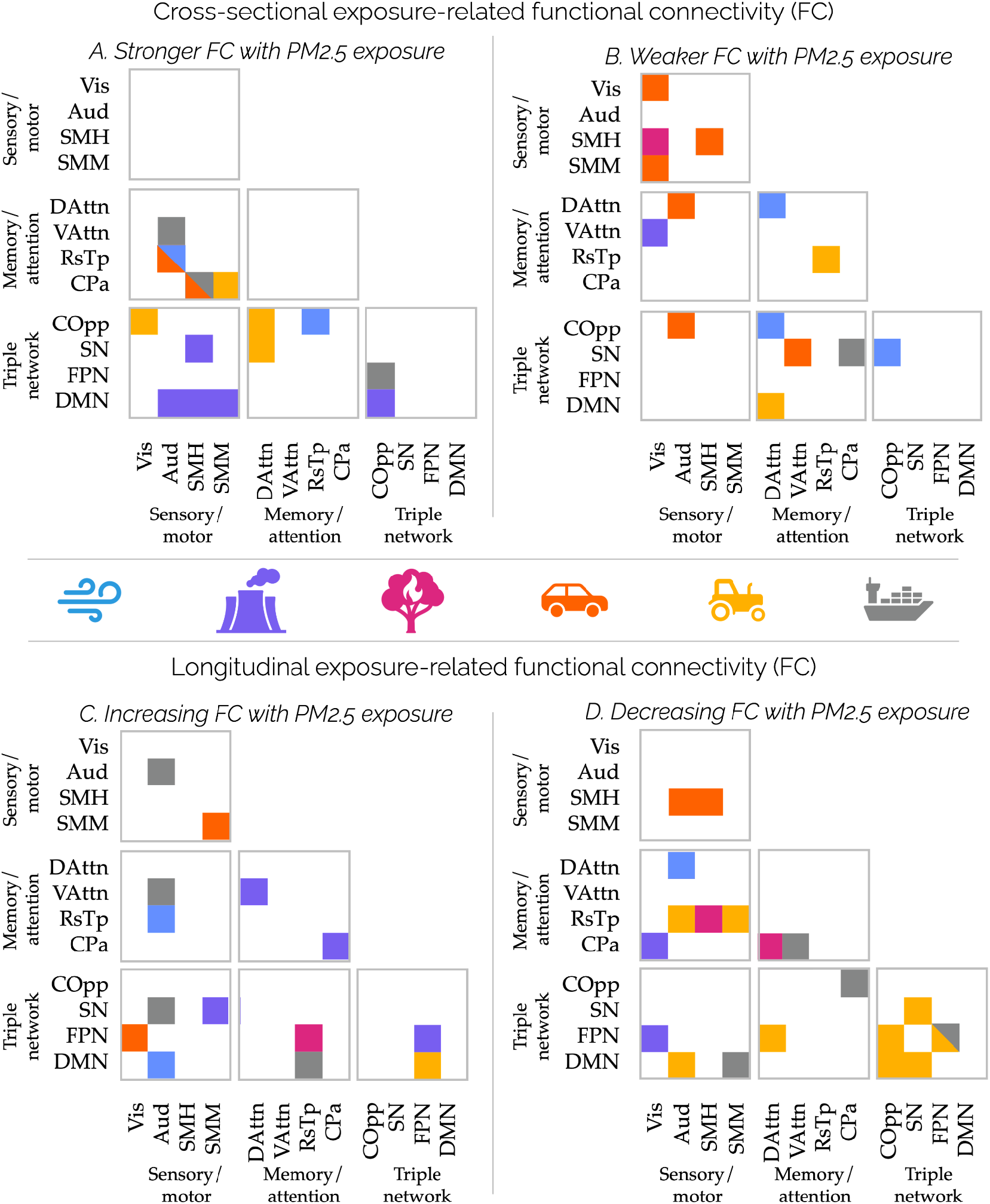
Summary of exposure-related connectivity by source, separated by direction of associations. Top: Cross-sectional functional connectivity positively (A) and negatively (B) related to exposure at ages 9-11 years, color-coded by source of PM_2.5_. Triangles indicate connections linked to multiple sources. Middle: Color-coded sources of PM_2.5_ (from left): crustal materials (blue), ammonium nitrates (yellow), biomass burning (pink), traffic (orange), ammonium sulfates (purple), and industrial/residual fuel burning (gray). Bottom: Longitudinal functional connectivity increasing (C) and decreasing (D) with greater exposure, from ages 9-13 years, color-coded by source of PM_2.5_. Triangles indicate connections linked to multiple sources. Networks are grouped into three overarching categories: sensory/motor networks (i.e., visual, Vis; auditory, Aud; somatosensory-hand, SMH; somatosensory-mouth, SMM), memory/attention networks (i.e., dorsal attention, DAttn; ventral attention, VAttn; retrosplenial temporal, RsTp; cinguloparietal, CPa), and triple network model (cingulo-opercular, COpp; salience, SN; frontoparietal, FPN; default mode, DMN).

### Geographical heterogeneity across ABCD Study sites

Comparing the predictive accuracy across the data collection sites (i.e., omitting one site for hardware-induced image artifacts) suggests some geographical heterogeneity in exposure-related rsFC and, potentially, differential generalizability of these findings across the U.S. (Table 2; Supplementary Table 5). For example, RMSE, which is on the same scale as the source exposure estimates, is similar in magnitude to site-wise standard deviations of exposure estimates (Supplementary Tables 3, 4).

**Table 2.** Fit of identified exposure-related FC across data collection sites.

| Source of PM<sub>2.5</sub> | Root mean squared error (RMSE) |  |
| --- | --- | --- |
|  | Differences at 9-10 years of age (N=20 sites) | Changes from 9-13 years of age (N=12 sites) |
| Crustal materials | 0.21 ± 0.11 | 0.21 ± 0.12 |
| Ammonium sulfates | 0.13 ± 0.05 | 0.13 ± 0.04 |
| Biomass burning | 0.12 ± 0.04 | 0.12 ± 0.03 |
| Traffic emissions | 0.20 ± 0.05 | 0.19 ± 0.05 |
| Ammonium nitrates | 0.17 ± 0.08 | 0.15 ± 0.07 |
| Industrial/residual fuel | 0.16 ± 0.04 | 0.16 ± 0.05 |

**Note.** Values represent the mean ± standard deviation across a leave-one-site-out cross-validation, using the CPM-identified pollution-related positive and negative connections as features in a linear regression to predict exposure per source (Equation 2). Root mean squared error indicates the gap between model-predicted and actual exposures, on the same scale as the exposure estimates (i.e., source contributions) themselves. Lower RMSE indicates a better performing model and, thus, a more generalizable pattern of exposure-related FC. See Supplementary Table 4 for $R^2$ per site.

That is, exposures predicted from rsFC are largely within one standard deviation of the true exposure estimates. Across sources of PM_2.5_, RMSE values suggest that the identified patterns of exposure-related rsFC best represent individuals in the northeast, but may not be representative of individuals from the west (Figure 6, Table 3). Overall, connectivity differences related to exposure to crustal materials and traffic showed the greatest heterogeneity across sites, while connectivity differences related to exposure to biomass burning and ammonium sulfates showed the least heterogeneity (Table 2, Supplementary Table 5). *Post hoc* correlations between RMSE and exposures by site (Supplementary Table 6), suggest that the identified patterns of exposure-related rsFC *differences* most poorly reflect individuals from sites with higher mean exposure to PM_2.5_ from crustal materials, biomass burning, and ammonium nitrates. Similarly, the identified exposure-related rsFC *changes* most poorly reflect individuals from sites with greater exposure to PM_2.5_ from crustal materials, traffic emissions, and ammonium nitrates. Further, the identified exposure-related rsFC *differences* and *changes* most poorly reflect individuals from sites with greater variability in exposure to PM_2.5_ from all sources (all *p* < 0.05). The proportion of variance in exposures explained by the identified patterns of positively- and negatively-related rsFC was small to negligible (i.e., per *R*^2^; Supplementary Table 5), suggesting that, after accounting for sources of bias in brain-exposure associations, PM_2.5_ exposure only plays a small role in rsFC differences between youth.

**Table 3.** Site RMSE averages by geographical region: Exposure-related connectivity at 9-11 years of age.

| Geographic<br>al Region<br>(Site N) | Crustal<br>material<br>s | Ammonium<br>sulfates | Biomass<br>burning | Traffic<br>emissions | Ammonium<br>nitrates | Industrial/<br>residual fuel | Mean |
| --- | --- | --- | --- | --- | --- | --- | --- |
| W (7) | 0.28 | 0.17 | 0.17 | 0.23 | 0.23 | 0.18 | 0.21 |
| SW (1) | 0.22 | 0.12 | 0.08 | 0.19 | 0.17 | 0.13 | 0.15 |
| MW (4) | 0.18 | 0.11 | 0.10 | 0.20 | 0.17 | 0.17 | 0.15 |
| NE (4) | 0.14 | 0.10 | 0.09 | 0.19 | 0.11 | 0.12 | 0.13 |
| SE (4) | 0.19 | 0.13 | 0.13 | 0.17 | 0.14 | 0.19 | 0.16 |
| Mean | 0.20 | 0.13 | 0.12 | 0.20 | 0.17 | 0.16 |  |

*Note.* Root mean squared error indicates the gap between model-predicted and actual exposures, on the same scale as the exposure estimates (i.e., source contributions) themselves. Lower RMSE indicates a better performing model and, thus, a more generalizable pattern of exposure-related FC. Geographical breakdown used to group study sites can be found in Supplementary Table 3. Abbreviations: W=west, SW=southwest, MW=midwest, NE=northeast, SE=southeast. For longitudinal models, not every geographical region was represented due to the reduced sample (i.e., only from sites with Siemens MRI scanners). RMSE for all cross-sectional and longitudinal exposure-related connectivity models is presented in Supplementary Table 4.

**Figure 6.**
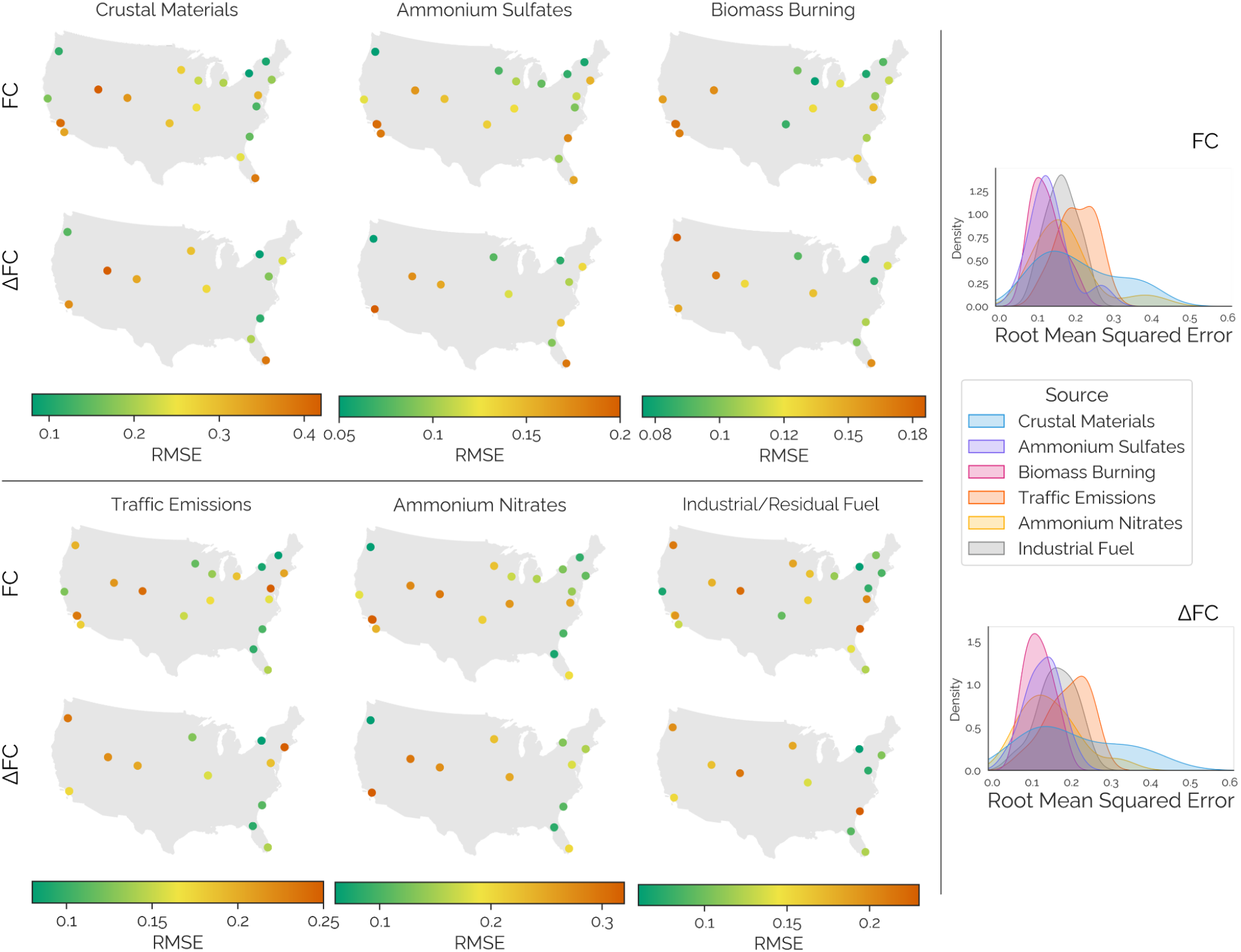
Geographical heterogeneity in the identified exposure-related FC patterns. Maps display model performance in a leave-one-site-out cross-validation scheme, as root mean squared error (RMSE) between actual and FC-predicted exposure for each source of PM_2.5_. Lower RMSE (green) indicates better model performance, while higher RMSE (red) indicates poorer model performance. For each plot, the colorbar limits are set by the distribution of RMSE per source, so that minima (i.e., green extrema) reflect the best-performing models while maxima (i.e., red extrema) reflect the poorest-performing models. Geographical heterogeneity of exposure-related FC patterns at ages 9-10 years (FC) in each of 20 sites (plotted by their latitude and longitude on a map of the United States), per source of PM_2.5_. Geographical generalizability of exposure-related FC changes (ΔFC) from age 9 to 13 years in each of 12 sites with Siemens MRI scanners (plotted by their latitude and longitude on a map of the United States), per source of PM_2.5_.

## Discussion

Here, we identified source-related exposure differences in the brain’s resting-state functional connectivity (rsFC) at ages 9-11 years and longitudinal changes in rsFC from age 9 to 13 years, using a large, geographically diverse U.S. cohort of children from the ABCD Study. Exposure-related differences in rsFC at ages 9-11 years partially overlapped with exposure-related changes in rsFC in the following two years. Sensorimotor networks showed the most exposure-related connectivity across sources, followed by the triple network model, while attention and memory networks exhibited the least. At ages 9-11 years, traffic emissions exposure was linked to a largely weaker connectivity of sensorimotor networks (Figure 5, top right), while ammonium sulfates and nitrates exposures were linked to largely stronger connectivity across the triple network model (Figure 5, top left). Exposure-related *changes* from 9-13 years were more widespread, suggesting that the transition to adolescence may be a time when the developing functional organization of the brain is particularly susceptible to sources of PM_2.5_ exposure. Exposure-related changes included decreasing connectivity linked to ammonium nitrate exposures (Figure 5, bottom right) and mixture of increasing and decreasing connectivity linked to industrial/residual fuel burning exposures (Figure 5, bottom). Geographically, the identified exposure-related connectivity patterns were more representative of youth from sites in the northeast and midwest, compared to youth from the southeast, west, and southwest. Altogether, opposing directions of exposure-related connectivity differences and changes are likely muddled in studies that estimate exposure from total PM_2.5_ and further complicated by geographical differences in PM_2.5_ sources.

### Sensorimotor networks related to sources of exposure at ages 9-11 years old

Theories of cognitive development suggest that sensorimotor functions are largely developed by childhood (Malik & Marwaha, 2025; Piaget, 1964). Developmental cognitive neuroscience, however, has shown that functional networks supporting these functions (e.g., visual, auditory, and somatomotor networks) continue to develop into early adolescence (Tooley et al., 2022), overlapping with higher-order networks that integrate sensorimotor information (e.g., default mode, salience, dorsal attention) and likely contributing to higher-order cognitive development (Bruchhage et al., 2020).

Here, the auditory network exhibited both stronger and weaker exposure-related connectivity at age 9-11 years, as well as exposure-related increases and decreases in connectivity from age 9-13 years, based on PM_2.5_ source and network. Due to the opposing directions, these exposure-related differences and changes are likely masked in studies that focus on total PM_2.5_. While the present work did not assess hearing or auditory processing, these findings contribute to a broader literature linking air pollution exposure to middle ear infections, hearing loss (both gradual and sudden), and central auditory system dysfunction (Gohari et al., 2024). Prior work has linked childhood middle ear infection incidence and hearing loss to exposure to metals and nitrates, in addition to PM_10_, PM_2.5_, and some gaseous pollutants (Calderón-Garcidueñas et al., 2011, 2017; Castellanos & Fuente, 2016; Gohari et al., 2024). Our findings extend this literature by showing that auditory network connectivity exhibited stronger connectivity at 9-11 years (e.g., with dorsal attention, retrosplenial temporal networks) and increasing connectivity at 9-13 years (e.g., with ventral attention, salience networks; within auditory network) with higher exposure to industrial/residual fuel emissions, which are high in metals. Further, studies of children who grew up in environments with extremely poor air quality show exposure-related lesions and conduction delays in auditory brainstem signals, and complementary postmortem histological studies showed that such children have lesions and neurodegenerative pathologies in auditory nuclei (Calderón-Garcidueñas et al., 2011). Likewise, we found weaker auditory network connectivity at ages 9-11 years (e.g., with dorsal attention, cingulo-opercular networks) with greater exposure to traffic emissions, and decreasing auditory network connectivity at 9-13 years (e.g., with retrosplenial temporal, default mode networks) with greater exposure to PM_2.5_ from traffic and ammonium nitrates. The retrosplenial temporal and default mode networks both support memory function (Kaboodvand et al., 2018; Santos et al., 2023), while the dorsal attention network is involved in top-down modulation of attention (Vossel et al., 2014). Thus, this work may present evidence of *in vivo* alterations to auditory functioning that accompany previously identified exposure-related harm to the auditory system and may suggest that such harms have further-reaching implications (e.g., for memory and attention). However, these results cannot indicate whether exposure-related connectivity differences are related to neurodegeneration in auditory nuclei, hearing loss, or infection.

Likewise, childhood connectivity of somatomotor networks differed based on exposure to PM_2.5_ from all sources assessed here, as did changes in somatomotor connectivity during the transition to adolescence. Unlike auditory network connectivity, however, somatomotor differences were more widespread at ages 9-11 years. Prior work has linked childhood motor impairment, both in humans and in animal models, to particulate air pollution from traffic, copper, and total PM_2.5_ (Costa et al., 2020; Lopuszanska & Samardakiewicz, 2020; Salvi & Salim, 2019) and, in this age range specifically, motor performance was poorer in children ages 8-12 years with greater environmental copper exposures (Pujol et al., 2016). Our findings may provide insight into the neural underpinnings of these exposure-related motor impairments. At 9-11 years of age, increased exposure to traffic and industrial/residual fuel emissions was linked to weaker somatomotor network connectivity with other sensory and motor networks, while greater exposure to biomass burning and traffic emissions was linked to weaker connectivity between somatomotor and visual networks. Altogether, this suggests childhood metal and carbon exposures may weaken the integration of sensory and motor information, impairing performance. Over the following two years, somatomotor network connectivity decreased with greater source-related exposure, including within the somatomotor-hand network (ammonium sulfates, traffic emissions, ammonium nitrates), with the retrosplenial temporal network (biomass burning, ammonium nitrates), and with the default mode network (industrial/residual fuel). Altogether, these findings suggest that exposure-related impairments in motor performance may be related to less coherent activity within somatomotor networks, to disrupted attention, or to multisensory integration (e.g., using visual or auditory information to direct movement). Further research should investigate exposure-related differences in brain function during a battery of sensorimotor and sensorimotor integration tasks to provide additional insight.

While the visual network exhibited fewer exposure-related differences in connectivity at ages 9-11 years and fewer exposure-related changes in connectivity over time from ages 9-13 years-old, there were a few notable trends. First, childhood visual network connectivity was weaker with greater exposure from ammonium sulfates, biomass burning, and traffic emissions, but stronger with greater exposure to ammonium nitrates, and largely unrelated to crustal and industrial/residual fuel exposure sources. During the transition to adolescence, longitudinal changes in visual network connectivity were negatively related to exposure, including connectivity with fronto-, cingulo-parietal networks with ammonium sulfates exposure, connectivity with somatomotor networks with biomass burning and within-visual network connectivity with traffic emissions. The only exception was positive associations between visual to somatomotor network connectivity seen with greater industrial/residential fuel exposure. Although the ABCD Study does not measure eye health, visual network differences may support a larger body of evidence linking PM_2.5_ exposure to eye health. In a large sample of adults, PM_2.5_ exposure was related to differences in retinal structure, which is supported by meta-analytic evidence of chronic eye diseases (Chua et al., 2020; Millen et al., 2024). Further meta-analytic evidence shows that PM_2.5_ exposure may have even greater effects on eye health during childhood and adolescence (Han et al., 2024). Additional bi-directional systems-level research is warranted to understand how pollutant driven changes in physical health may or may not contribute to brain-level differences in functional network organization and development.

### Triple network model is sensitive to secondary pollutants

Emotion regulation, social cognition, and executive control continue to develop throughout adolescence and into adulthood (National Academies of Sciences et al., 2019) and, when impaired, underlie many mental health conditions. The triple network model posits that aberrant connectivity between the default mode network, frontoparietal central executive network, and the salience network is responsible for cognitive dysfunction underlying multiple mental disorders (B. Menon, 2019; V. Menon, 2011). The default mode network, involved in self-referential processing and autobiographical memory, exhibits hyperconnectivity in depression (potentially contributing to rumination) but hypoconnectivity in autism (potentially underscoring social difficulties) (B. Menon, 2019). The frontoparietal central executive network, which maps on to the frontoparietal network delineation used here, supports working memory and goal-directed action in a range of cognitive tasks (V. Menon, 2011). Aberrant frontoparietal connectivity can include weaker within-network connectivity, compromising cognitive control; stronger connectivity between frontoparietal and regions of other networks; and weaker connectivity with the salience network, impairing access to relevant information. The salience network, which spans the salience and cingulo-opercular network delineations used here, detects and integrates both internal and external cues, filters “salient” information, and toggles between default mode and frontoparietal activity.

We found that salience, default mode, and, to a lesser extent, frontoparietal network connectivity with both sensorimotor and higher-order networks was associated with exposure to ammonium sulfates. Between higher-order networks, changes in connectivity were positively related to exposure (with the exception of negatively-related within-network changes in default mode connectivity). Prior work with these data, using a different methodology, found that, between default mode and salience networks, higher exposure (i.e., 8.68 µg/m^3^ annual average) was associated with increasing connectivity with age from 9 to 13 years, while lower exposure (6.66 µg/m^3^ annual average) was associated with decreasing connectivity with age (Cotter et al., 2023). Here, we linked decreasing exposure-related connectivity between default mode and cingulo-opercular networks with greater exposure to PM_2.5_ attributable to sources of ammonium nitrates, providing additional detail to these findings and adding evidence for exposure-related alterations in triple network connectivity. Between higher-order and sensorimotor networks, both differences in childhood connectivity and changes in connectivity during the transition to adolescence were positively associated with exposure. This may suggest that exposure to ammonium nitrates at ages 9-11 years may disrupt maturation of connectivity between the default mode and salience networks during early adolescence–potentially impacting the salience network’s ability to toggle between default mode and frontoparietal networks.

For ammonium nitrates, exposure-related differences in childhood connectivity in the triple network model largely involved the dorsal attention network, including weaker connectivity of default mode, but stronger connectivity of the salience network, with greater exposure. During the transition to adolescence, changes in triple network connectivity between the frontoparietal network and dorsal attention networks were positively related, while connectivity between the frontoparietal and default mode networks were negatively related, to ammonium nitrate exposure. Developmental trends in between-network connectivity suggest that higher-order networks (e.g., default mode, frontoparietal) become more segregated into adulthood (DeSerisy et al., 2021; Sherman et al., 2014). Thus, while we expect default mode and frontoparietal networks to be disconnected, in general, this negative exposure-related change in connectivity may potentially reflect accelerated development of these networks.

The triple network model suggests that unbalanced connectivity between default mode, salience and frontoparietal networks underlies psychopathology, and that dysfunction in any of the three networks may be responsible for related cognitive dysfunction (V. Menon, 2011). However, these three networks and the functional connectome at large are still developing in this age range. With these data it is difficult to determine whether the observed differences and changes in triple network connectivity reflect imbalance/dysfunction or transient alterations to developmental timing. Prior work assessing childhood PM_2.5_, NO_2_, and O_3_ exposure in this sample found no exposure-related increases in emotional problems from 9 to 13 years of age in the ABCD Study cohort (Campbell et al., 2023). However, more days with PM_2.5_ levels exceeding the EPA standard was linked to greater internalizing symptoms in the ABCD Study cohort (Smolker et al., 2024) and the broader literature includes widespread reports of poor mental health with greater exposure (Lian et al., 2023; Mota-Bertran et al., 2024; Newbury et al., 2021, 2024). This disconnect may be because the current sample spans from 9 to 13 years of age, and has not yet reached the peak age of mental health diagnoses, at 14.5 years (Solmi et al., 2022). If so, it is possible that these identified alterations to developing connectivity within and between these higher-order networks with transdiagnostic links to psychopathology may reflect early markers of eventual exposure-related mental health problems, or the subclinical changes preceding mental diagnoses. Future work should extend this research into mid-late adolescence to determine the potential clinical significance of these findings.

### Geographic heterogeneity in exposure-brain function associations and potential dose-dependent effects

A key aspect of this work is in comparing identified exposure-related rsFC patterns across multiple ABCD Study data collection sites, to examine geographical heterogeneity in these findings. Across study sites, exposure-related connectivity from biomass burning and ammonium sulfates were most generalizable. In contrast, exposure-related connectivity from crustal materials and traffic emissions were the most variable between sites. Across sources of PM_2.5_, exposure-related connectivity generalized least to youth living in the west United States but generalized most to youth living in the northeast. While cross-sectional model performance was significantly worse in sites with greater mean exposure to crustal, biomass burning, traffic emissions, and ammonium nitrates, longitudinal model performance was significantly related to mean exposure only for crustal materials, traffic emissions, and ammonium nitrates. Further, performance of all models (i.e., each source, cross-sectional and longitudinal) were related to sites’ standard deviations of exposure, potentially indicating greater heterogeneity in sites with more variable exposures between individuals. Together, this may indicate nonlinear or threshold effects of PM_2.5_ exposure on functional brain connectivity, such that brain-exposure associations at low PM_2.5_ concentrations poorly reflect those observed at higher concentrations. Additional research should aim to better characterize brain-exposure associations across a wider range of PM_2.5_ exposure, as this sample’s average exposure was a whole standard deviation below the EPA standard (9.0 µg/m^3^ annual average).

Additionally, it is possible that differences in participant demographics between regions and sites play a role in the predictability of these models, as predictive models perform worse on individuals who do not conform to sample stereotypes (Greene et al., 2022). Although the ABCD Study sample is large and diverse, youth from lower income, less educated, and non-white households are underrepresented, compared to the national demographic composition and varying demographic breakdowns of each data collection site. Here, we see poor generalizability of exposure-related rsFC to sites in the west and southeast, and in sites with more variable exposures. Additional research should explain these discrepancies in geographical generalizability and whether they are related to dose-dependent impacts of exposure or differences in demographic composition of samples from each site. Further, differences in generalizability of the identified brain-exposure phenomena due to geography and due to the different MRI scanners used to acquire the neuroimaging data used here are inextricable, as sites use different scanners from only one of three manufacturers (i.e., SEMENS, General Electric, and Philips). While there exist many methods of data harmonization that address between-manufacturer and between-scanner differences in MRI data, they are ill-suited to predictive modeling due to data leakage they introduce between model training and testing steps (Rosenblatt et al., 2024). Future work should aim to quantify and mitigate these differences in predictive modeling frameworks for more robust multi-dataset investigations, which greatly increase the statistical power and generalizability of brain-exposure and brain-phenotype investigations.

### Strengths & Limitations

In addition to the aforementioned strengths and limitations incurred by the ABCD Study multi-site design, the modeling approaches used here confer both strengths and limitations to this work. First, in the rsFC literature, researchers are coming to value prediction over reliability in phenotype- and, in this case, exposure-related brain networks because they are more generalizable and applicable in real-world settings (Finn & Rosenberg, 2021). Second, the rsFC estimates used here were computed at the network level, instead of at a finer-grained region level. Strengths of this approach are that these networks represent canonical cortical subdivisions (Gordon et al., 2016, 2017; Power et al., 2011; Uddin et al., 2019), that signal-to-noise ratio is higher at the network-level (i.e., by implicitly applying spatial smoothing), and that statistical power is higher due to theory-driven dimensionality reduction. To this last point, the network-level rsFC estimates used here come from 12 large-scale brain networks, summarizing rsFC of 300 brain regions. Because connectivity estimates are pairwise, this reduces the number of unique connections analyzed from 44,850 to 78. Weaknesses of assessing network-level, instead of regional, brain connectivity are that they overlook finer-grained specializations of their comprising regions and connections, that they may wash out relevant subnetwork dynamics (e.g., default mode network subdivisions have exhibited functional specializations (Laird et al., 2009)), and that they may be less robust than finer-grained connectivity estimates (Noble et al., 2017; J. Wang et al., 2017).

## Conclusions

Here, we identified and assessed patterns of exposure-related childhood rsFC at ages 9-11 years and changes during the transition to adolescence, from age 9-13 years, with respect to several sources of outdoor PM_2.5_. Across these patterns, we identified several themes in exposure-related connectivity of sensorimotor networks and the triple network model of psychopathology. Exposure-related differences in visual, auditory, and somatomotor network connectivity were notably related to sources high in metals (e.g., traffic emissions, industrial/residential fuels) and ammonium nitrates. These findings may provide insight into exposure-related auditory, visual, and motor deficiencies noted previously in epidemiological research, their neural correlates, and their implications for other brain functions. Prior work suggests that children may be more susceptible to such deficiencies and, as the functional connectome is still developing, these findings may show how childhood sensory and motor detriments can propagate into altered connectivity of other networks and, thus, alterations of other functions. On the other hand, connectivity of the “triple network model”, in which connectivity imbalances are broadly related to mental health conditions, was largely related to secondary sources of PM_2.5_ (i.e., ammonium sulfates and nitrates). Together with prior findings, the exposure-related connectivity between default mode, salience, and frontoparietal networks may be early markers of exposure-related mental health conditions, which likely arise later in adolescence. The geographical generalizability of these patterns of exposure-related rsFC may indicate dose-dependent effects of PM_2.5_ from crustal materials, biomass burning, and ammonium nitrates, but further research is needed. Together, these findings may provide insight into some exposure-related sensory and motor difficulties observed in epidemiological studies, while providing insight into potential early markers of exposure-related mental health problems that may arise later in adolescence. Additional longitudinal neurodevelopment research is needed to better understand and support such biomarkers.

## Acknowledgment

A special thank you to all of the children and families for their participation in their ABCD Study.

Research described in this article was supported by the National Institutes of Health (MMH: NIEHS R01ES032295, R01ES031074; KLB: P30ES07048-27) CCI would like to acknowledge scholars involved in NSP (R25 NS089462), BRAINS (R25 NS094094), and Diversifying CNS (R25 NS117356), as well as R25MH125545 and R25MH120869 for creating a supportive network of ABCD Study users.

Data used in the preparation of this article were obtained from the Adolescent Brain Cognitive Development^SM^ (ABCD) Study (https://abcdstudy.org), held in the NIMH Data Archive (NDA). This is a multisite, longitudinal study designed to recruit more than 10,000 children aged 9-11 years old and follow them over 10 years into early adulthood. The ABCD Study® is supported by the National Institutes of Health and additional federal partners under award numbers U01DA041048, U01DA050989, U01DA051016, U01DA041022, U01DA051018, U01DA051037, U01DA050987, U01DA041174, U01DA041106, U01DA041117, U01DA041028, U01DA041134, U01DA050988, U01DA051039, U01DA041156, U01DA041025, U01DA041120, U01DA051038, U01DA041148, U01DA041093, U01DA041089, U24DA041123, U24DA041147. A full list of supporters is available at https://abcdstudy.org/federal-partners.html. A listing of participating sites and a complete listing of the study investigators can be found at https://abcdstudy.org/consortium_members/. ABCD consortium investigators designed and implemented the study and/or provided data but did not necessarily participate in the analysis or writing of this report. This manuscript reflects the views of the authors and may not reflect the opinions or views of the NIH or ABCD consortium investigators. The ABCD data repository grows and changes over time. The ABCD data used in this report came from https://dx.doi.org/10.15154/z563-zd24.

## Author Contributions

Conceptualization: KLB, MMH

Data curation: KLB, KS, JS, AVDJ

Formal Analysis: KLB, KS

Funding acquisition: MMH, KLB

Methodology: KLB, MMH, RH, DAH

Project administration: KLB, MMH

Resources: MMH, JS

Software: KLB, KS

Supervision: MMH

Visualization: KLB

Writing – original draft: KLB, MMH

Writing – review & editing: KLB, KS, RH, JS, MMH, DAH

