## Supplementary Information for "Sources of outdoor air pollution exposure and child brain network development across the United States"

### Outdoor air pollution exposure and functional network development in children: Source-specific predictive connectomics

This document includes:

- Supplementary Methods
- Supplementary Tables 1 to 6
- Supplementary Figures 1 to 4
- Supplementary References

*Corresponding Author*: **Dr. Megan Herting**, Associate Professor, Department of Population and Public Health Sciences, Keck School of Medicine, University of Southern California, Los Angeles, CA USA |

### Supplementary Methods

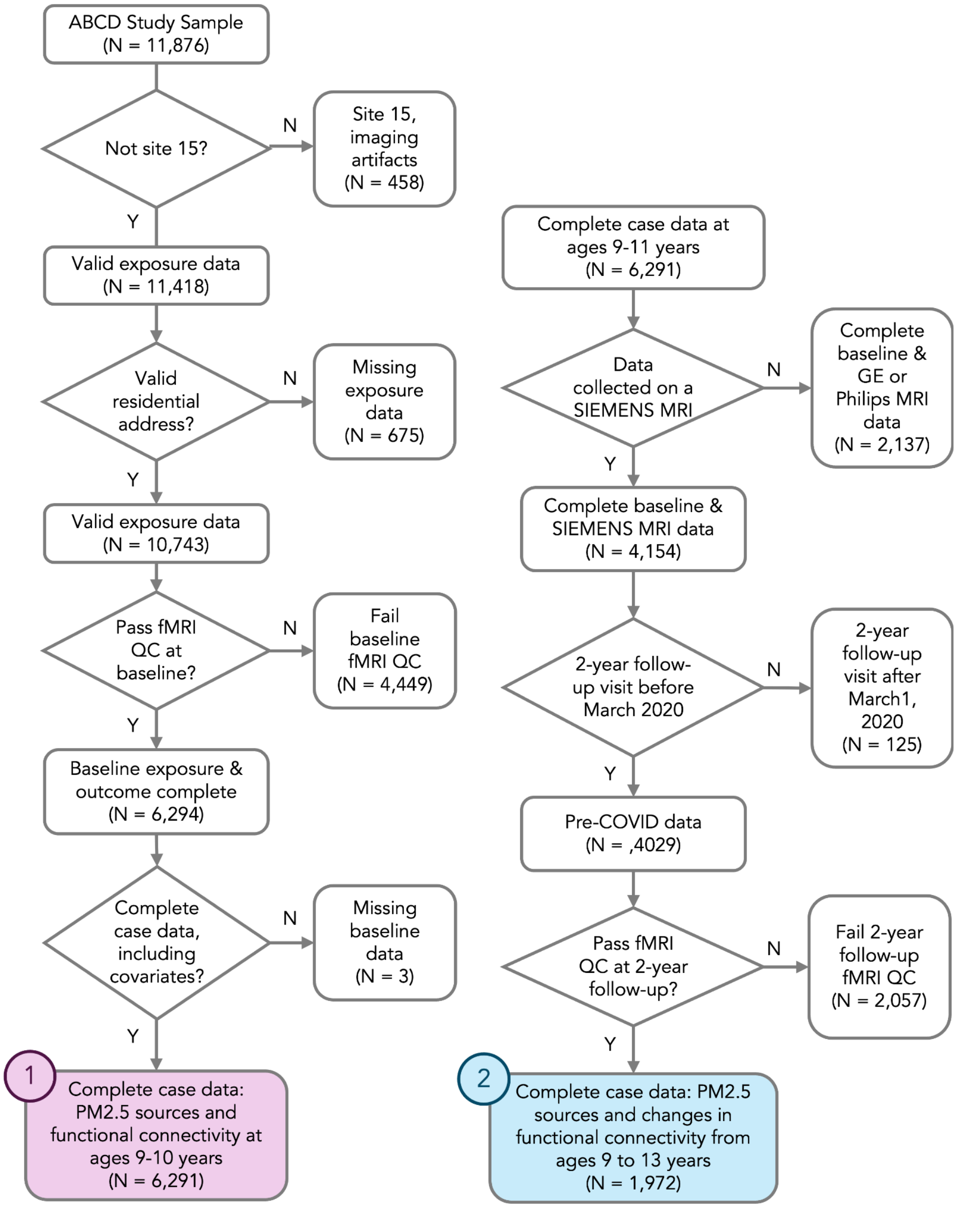


***Supplementary Figure 1. Participant inclusion and exclusion across criteria.*** (1) indicates the final cross-sectional sample size and (2) indicates the final longitudinal sample size.

#### Directed Acyclic Graph (DAG)

Using the notion that a confounder must be related to both the exposure (X) and the outcome (Y) (Shrier & Platt, 2008), we used the software DAGgitty (Textor et al., 2016) to assist in the creation and interpretation of a DAG for each specific aim of our study. Using color-coded variables and unidirectional arrows we indicate how potential confounding and precision variables are interrelated and their relations to both exposure and outcome, to identify biasing pathways. This graph (Supplementary Figure 1) was completed using field-specific background knowledge, familiarity with published literature, and intuition from authors MMH, JC, and JS. Since exposure estimates are based on the child's primary residential address, a potential confounder (c) must predict both the child's residential location (X) and their brain outcome (Y). Using the below causal diagram and the software DAGgitty, we identified and reviewed potential biasing and backdoor (i.e., biasing, noncausal correlations between X and Y) paths among these variables to identify minimal adjustment sets to reduce potential confounding.

The minimal adjustment sets identified by our DAG analysis, along with MRI precision variables, are outlined in bold in Supplementary Figure 2. Confounding variables include and include critical confounders such as youth race/ethnicity, sex, and age (yellow); parental sociodemographic factors such as household income (green); other environmental factors such as perceived neighborhood safety, population density, distance to major roadways, and urban vs. rural classification of their primary residential location (purple); and variables describing participant engagement in outdoor environment such as average weekly physical activity and average screen time use (orange). Additionally, precision variables such as MRI manufacturer and head motion during MRI scans (blue) were added, in addition to youth handedness. See Morrel et al., for a review of outdoor air pollution influences on child and adolescent brain development (Morrel et al., 2025) including confounding and precision variables commonly used in such studies.

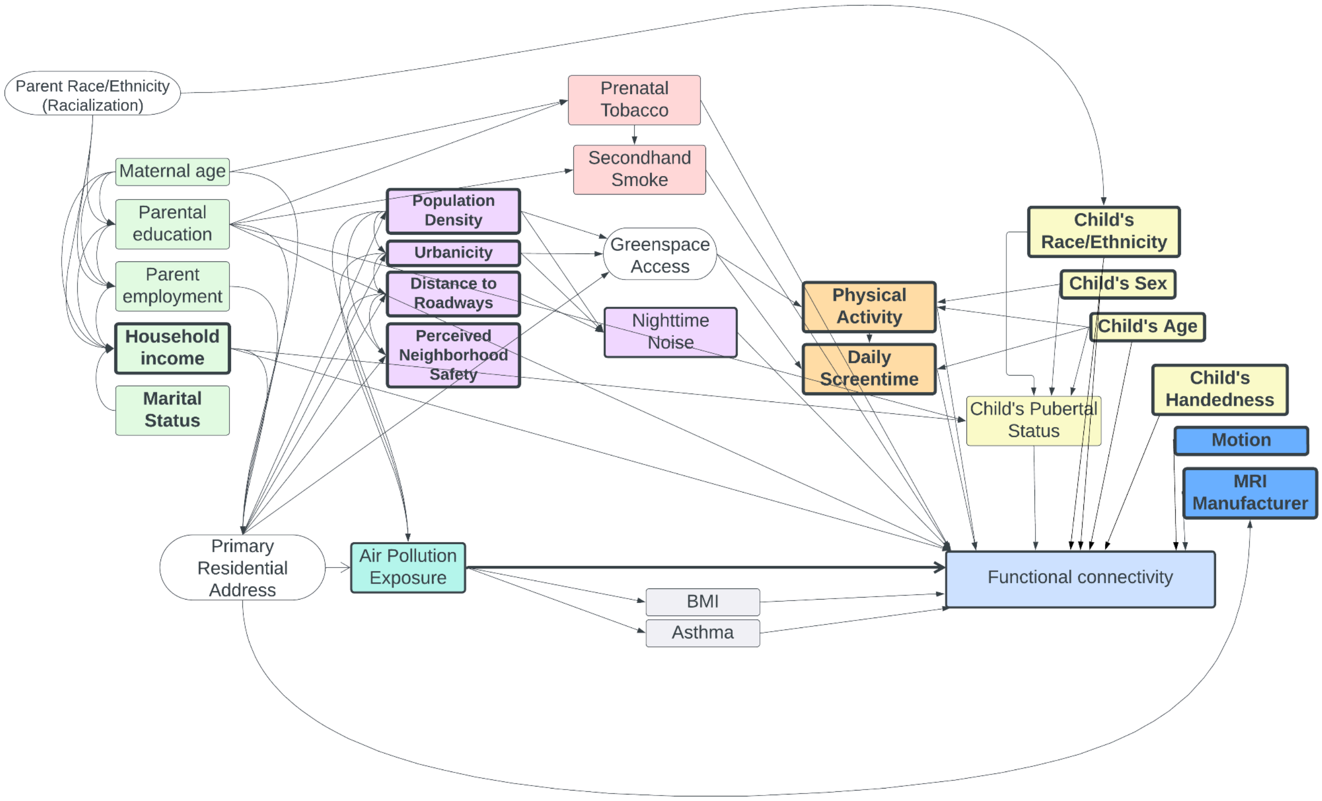


***Supplementary Figure 2***. Directed acyclic graph of factors contributing to air pollution exposure and changes in functional connectivity. Potential confounding variables are color-coded according to the following categories: parental sociodemographic factors (green), other environmental factors (purple), pregnancy-related factors (pink), participant engagement in outdoor environment (orange), participant characteristics yellow), precision MRI variables (dark blue), and health variables gray). All rectangular boxes represent measured variables, while rounded/oval boxes represent unmeasured constructs. Each arrow indicates a causal relationship between two variables. Thick outlines represent variables included in the "minimally sufficient adjustment set" (or vital precision variables) that were regressed out of exposure and connectivity measures before predictive models were run.

***Supplementary Table 1.*** Performance of models predicting PM2.5 component concentrations as reported by Amini et al. (2021).

| **Component** | **Model RMSE** |
|---|---|
| EC | 0.09 |
| $NH_4^+$ | 0.09 |
| $NO_3^-$ | 0.07 |
| OC | 0.18 |
| $SO_4^{2-}$ | 0.28 |
| Br | 0.18 |
| Ca | 2.86 |
| Cu | 1.35 |
| Fe | 4.99 |
| K | 4.77 |
| Ni | 0.45 |
| Pb | 0.34 |
| Si | 10.15 |
| V | 0.73 |
| Zn | 1.82 |

*Abbreviations*: nickel, Ni; iron, Fe; bromine, Br; vanadium, V; potassium, K; lead, Pb; zinc, Zn; copper, Cu; calcium, Ca; silicon, Si; elemental carbon, EC; sulfates, $SO_4^{2-}$; nitrates, $NO_3^-$; ammonium, $NH_4^+$; and organic carbon, OC

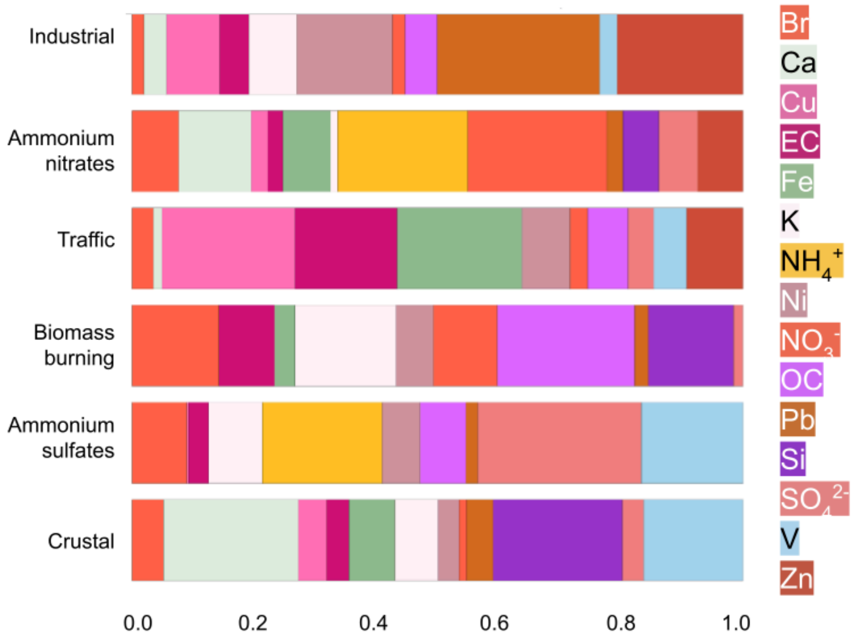


***Supplementary Figure 3.*** Loadings of each $PM_{2.5}$ component on PMF-derived common sources of outdoor air pollution for all ABCD study participants across the 21 study sites. Adapted from Sukumaran et al., 2024 & Bottenhorn et al., 2024.

### Supplementary Results

***Supplementary Table 2.*** Descriptive statistics of each chemical component of $PM_{2.5}$

| | Cross-sectional sample (*N* = 6291) | | Longitudinal sample (*N* = 1972) | |
|---|---|---|---|---|
| | Mean | Standard deviation | Mean | Standard deviation |
| | | ng/m³ | | |
| Br | 2.58 | 0.57 | 2.60 | 0.54 |
| Ca | 50.40 | 21.71 | 51.59 | 22.01 |
| Cu | 4.53 | 1.58 | 4.44 | 1.47 |
| Fe | 65.54 | 23.42 | 63.82 | 20.99 |
| K | 62.86 | 9.66 | 61.94 | 8.64 |
| Ni | 0.82 | 0.25 | 0.79 | 0.23 |
| Pb | 4.46 | 1.13 | 4.47 | 1.18 |
| Si | 88.62 | 39.54 | 93.42 | 42.16 |
| V | 0.37 | 0.20 | 0.41 | 0.21 |
| Zn | 8.86 | 3.36 | 8.60 | 3.10 |
| | | μg/m³ | | |
| EC | 0.52 | 0.15 | 0.51 | 0.13 |
| $NH_4^+$ | 0.29 | 0.12 | 0.25 | 0.11 |
| $NO_3^-$ | 0.94 | 0.34 | 0.90 | 0.33 |
| OC | 1.86 | 0.44 | 1.76 | 0.41 |
| $SO_4^{2-}$ | 0.88 | 0.28 | 0.80 | 0.28 |

*Abbreviations*: nickel, Ni; iron, Fe; bromine, Br; vanadium, V; potassium, K; lead, Pb; zinc, Zn; copper, Cu; calcium, Ca; silicon, Si; elemental carbon, EC; sulfates, $SO_4^{2-}$; nitrates, $NO_3^-$; ammonium, $NH_4^+$; and organic carbon, OC

***Supplementary Table 3***. Average and standard deviation of participant contributions to each of the six common $PM_{2.5}$ sources for each data collection site, cross-sectional sample.

| | *N* | Crustal | Ammonium Sulfates | Biomass Burning | Traffic | Ammonium Nitrates | Industrial/ Residual Fuel |
|---|---|---|---|---|---|---|---|
| *West* | | | | | | | |
| CHLA | 406 | 0.99 ± 0.36 | 1.28 ± 0.29 | 1.63 ± 0.22 | 2.03 ± 0.36 | 2.02 ± 0.51 | 0.72 ± 0.22 |
| CUB | 558 | 1.82 ± 0.35 | 0.11 ± 0.16 | 1.11 ± 0.14 | 1.01 ± 0.4 | 0.89 ± 0.27 | 0.66 ± 0.28 |
| UTAH | 1012 | 1.87 ± 0.42 | 0.03 ± 0.15 | 1.29 ± 0.17 | 0.88 ± 0.29 | 1.45 ± 0.23 | 0.98 ± 0.22 |
| SRI | 350 | 0.53 ± 0.14 | 0.64 ± 0.14 | 1.43 ± 0.16 | 1.32 ± 0.23 | 0.85 ± 0.22 | 1.0 ± 0.13 |
| UCLA | 434 | 1.1 ± 0.37 | 1.22 ± 0.28 | 1.64 ± 0.22 | 1.77 ± 0.29 | 1.56 ± 0.43 | 0.73 ± 0.18 |
| UCSD | 737 | 1.22 ± 0.26 | 0.96 ± 0.18 | 1.36 ± 0.18 | 1.41 ± 0.31 | 1.07 ± 0.19 | 0.93 ± 0.2 |
| OHSU | 583 | 0.34 ± 0.14 | 0.44 ± 0.08 | 1.48 ± 0.2 | 1.19 ± 0.33 | 0.23 ± 0.11 | 1.25 ± 0.32 |
| *Southwest* | | | | | | | |
| LIBR | 744 | 2.0 ± 0.27 | 1.36 ± 0.15 | 0.87 ± 0.09 | 0.73 ± 0.26 | 0.81 ± 0.21 | 0.87 ± 0.15 |
| *Midwest* | | | | | | | |
| UMICH | 728 | 0.15 ± 0.19 | 1.1 ± 0.13 | 0.78 ± 0.12 | 0.8 ± 0.42 | 1.75 ± 0.18 | 1.26 ± 0.2 |
| UMN | 602 | 0.38 ± 0.26 | 0.75 ± 0.11 | 0.83 ± 0.11 | 0.55 ± 0.34 | 1.47 ± 0.18 | 1.17 ± 0.31 |
| UWM | 384 | 0.31 ± 0.2 | 0.9 ± 0.12 | 0.7 ± 0.08 | 0.97 ± 0.26 | 1.77 ± 0.16 | 1.52 ± 0.19 |
| WUSTL | 707 | 1.1 ± 0.21 | 1.33 ± 0.15 | 0.73 ± 0.14 | 0.79 ± 0.41 | 1.25 ± 0.19 | 1.16 ± 0.29 |
| *Northeast* | | | | | | | |
| ROC | 339 | 0.14 ± 0.1 | 1.12 ± 0.09 | 0.7 ± 0.1 | 0.59 ± 0.16 | 1.24 ± 0.15 | 1.15 ± 0.13 |
| UMB | 604 | 0.41 ± 0.25 | 1.37 ± 0.15 | 0.74 ± 0.11 | 1.29 ± 0.43 | 0.95 ± 0.15 | 1.27 ± 0.16 |
| UVM | 579 | 0.31 ± 0.17 | 0.79 ± 0.1 | 0.92 ± 0.11 | 0.23 ± 0.2 | 0.53 ± 0.13 | 0.68 ± 0.26 |
| YALE | 600 | 0.42 ± 0.14 | 1.11 ± 0.18 | 0.8 ± 0.12 | 0.98 ± 0.43 | 0.63 ± 0.15 | 1.33 ± 0.15 |
| *Southeast* | | | | | | | |
| MUSC | 378 | 1.12 ± 0.14 | 1.54 ± 0.17 | 1.16 ± 0.11 | 0.74 ± 0.21 | -0.05 ± 0.11 | 0.81 ± 0.28 |
| FIU | 631 | 2.34 ± 0.43 | 1.31 ± 0.16 | 0.41 ± 0.15 | 1.16 ± 0.22 | 0.39 ± 0.18 | 0.47 ± 0.16 |
| UFL | 448 | 1.75 ± 0.19 | 1.49 ± 0.14 | 1.2 ± 0.15 | 0.61 ± 0.22 | -0.05 ± 0.13 | 0.23 ± 0.24 |
| VCU | 550 | 0.46 ± 0.13 | 1.51 ± 0.17 | 0.96 ± 0.16 | 0.83 ± 0.35 | 0.59 ± 0.21 | 0.95 ± 0.32 |

*Abbreviations*. CHLA, Children's Hospital of Los Angeles; CUB, University of Colorado Boulder; FIU, Florida

International University; LIBR, Laureate Institute for Brain Research; MUSC, Medical University of South Carolina; OHSU, Oregon Health and Science University; ROC, University of Rochester; SRI, SRI International; UCLA, University of California, Los Angeles; UCSD, UC San Diego; UFL, University of Florida; UMB, University of Maryland Baltimore; UMICH, University of Michigan; UMN, University of Minnesota; UTAH, University of Utah; UVM, University of Vermont; UWM, University of Wisconsin—Milwaukee; VCU, Virginia Commonwealth University; WUSTL, Washington University in St. Louis; YALE, Yale University.

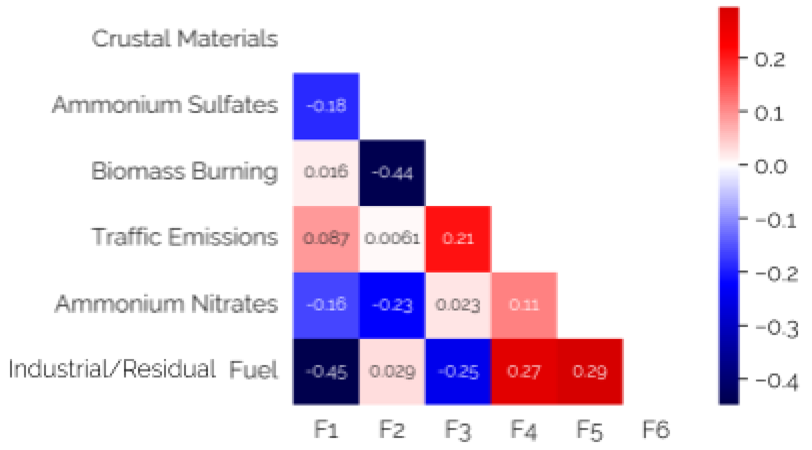


***Supplementary Figure 4***. Correlations between loadings on each PMF-derived source of $PM_{2.5}$.

***Supplementary Table 4.*** RMSE per site for connectivity-predicted exposure for each source.

| Site | Crustal | | Ammonium Sulfates | | Biomass | | Traffic | | Ammonium Nitrates | | Industrial/ Residual Fuel | |
|---|---|---|---|---|---|---|---|---|---|---|---|---|
| | 9-11 years | 9-13 years | 9-11 years | 9-13 years | 9-11 years | 9-13 years | 9-11 years | 9-13 years | 9-11 years | 9-13 years | 9-11 years | 9-13 years |
| *West* | | | | | | | | | | | | |
| CHLA | 0.38 | – | 0.27 | – | 0.19 | – | 0.25 | – | 0.41 | – | 0.20 | – |
| CUB | 0.32 | 0.31 | 0.13 | 0.15 | NaN | 0.12 | 0.26 | 0.24 | 0.24 | 0.20 | 0.22 | 0.21 |
| UTAH | 0.40 | 0.42 | 0.16 | 0.16 | 0.17 | 0.16 | 0.25 | 0.24 | 0.20 | 0.21 | 0.18 | 0.19 |
| SRI | 0.13 | – | 0.12 | – | 0.14 | – | 0.17 | – | 0.17 | – | 0.11 | – |
| UCLA | 0.34 | 0.32 | 0.27 | 0.20 | 0.21 | 0.14 | 0.25 | 0.21 | 0.35 | 0.32 | 0.17 | 0.16 |
| UCSD | 0.27 | – | 0.17 | – | 0.17 | – | 0.22 | – | 0.18 | – | 0.16 | – |
| OHSU | 0.10 | 0.09 | 0.05 | 0.05 | NaN | 0.18 | 0.24 | 0.24 | 0.08 | 0.06 | 0.22 | 0.21 |
| *Southwest* | | | | | | | | | | | | |
| LIBR | 0.22 | – | 0.12 | – | 0.08 | – | 0.19 | – | 0.17 | – | 0.13 | – |
| *Midwest* | | | | | | | | | | | | |
| UMICH | 0.15 | – | 0.11 | – | 0.11 | – | 0.24 | – | 0.15 | – | 0.14 | – |
| UMN | 0.22 | 0.23 | 0.09 | 0.09 | 0.09 | 0.08 | 0.17 | 0.16 | 0.18 | 0.16 | 0.19 | 0.19 |
| UWM | 0.16 | – | 0.11 | – | 0.08 | – | 0.18 | – | 0.16 | – | 0.17 | – |
| WUSTL | 0.20 | 0.18 | 0.12 | 0.13 | 0.13 | 0.13 | 0.20 | 0.19 | 0.19 | 0.18 | 0.17 | 0.16 |
| *Northeast* | | | | | | | | | | | | |
| ROC | 0.08 | 0.08 | 0.09 | 0.09 | 0.08 | 0.07 | 0.12 | 0.08 | 0.11 | 0.11 | 0.09 | 0.06 |
| UMB | 0.25 | 0.13 | 0.12 | 0.11 | 0.09 | 0.08 | 0.29 | 0.24 | 0.13 | 0.13 | 0.11 | 0.12 |
| UVM | 0.09 | – | 0.07 | – | 0.09 | – | 0.11 | – | 0.10 | – | 0.14 | – |
| YALE | 0.14 | 0.16 | 0.14 | 0.14 | 0.11 | 0.10 | 0.25 | 0.25 | 0.11 | 0.11 | 0.12 | 0.13 |
| *Southeast* | | | | | | | | | | | | |
| MUSC | 0.12 | 0.08 | 0.16 | 0.15 | 0.10 | 0.10 | 0.16 | 0.16 | 0.10 | 0.08 | 0.24 | 0.23 |
| FIU | 0.38 | 0.37 | 0.15 | 0.17 | 0.14 | 0.14 | 0.19 | 0.17 | 0.17 | 0.13 | 0.15 | 0.15 |
| UFL | 0.17 | 0.15 | 0.11 | 0.10 | 0.13 | 0.10 | 0.15 | 0.13 | 0.09 | 0.07 | 0.16 | 0.13 |
| VCU | 0.10 | – | 0.11 | – | 0.14 | – | 0.20 | – | 0.19 | – | 0.21 | – |

*Abbreviations*. CHLA, Children's Hospital of Los Angeles; CUB, University of Colorado Boulder; FIU, Florida International University; LIBR, Laureate Institute for Brain Research; MUSC, Medical University of South Carolina; OHSU, Oregon Health and Science University; ROC, University of Rochester; SRI, SRI International; UCLA, University of California, Los Angeles; UCSD, UC San Diego; UFL, University of Florida; UMB, University of Maryland Baltimore; UMICH, University of Michigan; UMN, University of Minnesota; UTAH, University of Utah; UVM, University of Vermont; UWM, University of Wisconsin—Milwaukee; VCU, Virginia Commonwealth University; WUSTL, Washington University in St. Louis; YALE, Yale University.

***Supplementary Table 5.*** Coefficient of determination between actual and predicted exposure to each source of $PM_{2.5}$, from leave-one-site-out cross-validation.

| Pollution source | Ages 9-10 years N=20 sites | | Ages 9-13 years N=12 sites | |
|---|---|---|---|---|
| | Mean $R^2$ | Standard deviation | Mean $R^2$ | Standard deviation |
| Crustal | -0.011 | 0.018 | -0.017 | 0.032 |
| Ammonium nitrates | -0.004 | 0.011 | -0.024 | 0.045 |
| Biomass | -0.001 | 0.006 | -0.012 | 0.017 |
| Traffic | -0.008 | 0.009 | 0.000 | 0.018 |
| Ammonium sulfates | -0.008 | 0.018 | -0.023 | 0.041 |
| Industrial/residual fuel | -0.003 | 0.011 | 0.002 | 0.033 |

*Note.* Coefficient of determination, $R^2$ indicates the proportion of variance in exposure explained by the identified exposure-related FC patterns. Higher numbers indicate a better performing model and, thus, a more generalizable pattern of exposure-related FC, while negative numbers indicate how much worse the trained model performed compared to an intercept-only model.

***Supplementary Table 6.*** *Post hoc* correlations between site RMSE and both mean and standard deviation of exposure per source of $PM_{2.5}$, for cross-sectional and longitudinal models

| | | Mean Exposure, by Site | | Standard Deviation of Exposure, by Site | |
|---|---|---|---|---|---|
| | | *r* | *p* | *r* | *p* |
| Crustal materials | Cross-sectional | **0.642** | **0.002** | **0.862** | **0.000** |
| | Longitudinal | *0.629* | *0.028* | **0.972** | **0.000** |
| Ammonium sulfates | Cross-sectional | 0.195 | 0.409 | **0.911** | **0.000** |
| | Longitudinal | 0.028 | 0.931 | **0.881** | **0.000** |
| Biomass burning | Cross-sectional | **0.662** | **0.003** | **0.930** | **0.000** |
| | Longitudinal | 0.371 | 0.236 | **0.944** | **0.000** |
| Traffic emissions | Cross-sectional | **0.681** | **0.001** | **0.650** | **0.002** |
| | Longitudinal | *0.629* | *0.028* | **0.713** | **0.009** |
| Ammonium nitrates | Cross-sectional | **0.617** | **0.004** | **0.941** | **0.000** |
| | Longitudinal | **0.818** | **0.001** | **0.993** | **0.000** |
| Industrial /residual fuel | Cross-sectional | -0.209 | 0.376 | **0.758** | **0.000** |
| | Longitudinal | -0.077 | 0.812 | *0.650* | *0.022* |

***Note.*** **Bold** values indicate significant correlation between model performance and site exposure mean/variability at $\alpha < 0.01$, *italicized* values indicate significant correlation between model performance and site exposure mean/variability at $\alpha < 0.05$. "Cross-sectional" models estimate exposure-related differences in functional connectivity at ages 9-10 years, while "longitudinal" models estimate exposure-related changes in functional connectivity from 9 to 13 years of age but only in sites with Siemens MRI scanners.
